# AI-Driven Knowledge Synthesis for Food Web Parameterisation

**DOI:** 10.1101/2025.05.18.654761

**Authors:** Scott Spillias, Elizabeth A. Fulton, Fabio Boschetti, Cathy Bulman, Joanna Strzelecki, Rowan Trebilco

## Abstract

We introduce a proof-of-concept framework, Synthesising Parameters for Ecosystem modelling with LLMs (SPELL), that automates species grouping and diet matrix generation to accelerate food web construction for ecosystem models. SPELL retrieves species lists, classifies them into functional groups, and synthesizes trophic interactions by integrating global biodiversity databases (e.g., FishBase, GLOBI), species interaction repositories, and optionally curated local knowledge using Large Language Models (LLMs). We validate the approach through a marine case study across four Australian regions, achieving high reproducibility in species grouping (>99.7%) and moderate consistency in trophic interactions (51-59%). Comparison with an expert-derived food web for the Great Australian Bight indicates strong but incomplete ecological accuracy: 92.6% of group assignments were at least partially correct and 82% of trophic links were identified. Specialized groups such as benthic organisms, parasites, and taxa with variable feeding strategies remain challenging. These findings highlight the importance of expert review for fine-scale accuracy and suggest SPELL is a generalizable tool for rapid prototyping of trophic structures in marine and potentially non-marine ecosystems.

**Highlights:**

- LLM-based framework automates species grouping and diet matrix creation with >99.7% consistency
- 51–59% of trophic interactions show high stability (stability score > 0.7) across iterations
- In expert comparison, SPELL achieved 81.6% agreement and 80% of diet differences < 0.2
- LLM-driven synthesis integrates global databases with unstructured local knowledge
- Reduces ecosystem model development time from months to hours

## 1. Introduction

Ecosystem modelling is an important tool for understanding and managing complex environments, with food web models being essential for representing trophic interactions and energy flows in marine ecosystems and predicting their responses to external pressures [8, 11, 2]. These models provide quantitative insights into ecosystem structure and function, enabling researchers to assess cumulative impacts of multiple stressors and support ecosystem-based management decisions [10, 43, 16]. However, constructing these models presents significant challenges, particularly in developing food webs that capture the complex web of trophic interactions within an ecosystem.

Traditional approaches to food web construction rely heavily on extensive literature review, data collation and expert knowledge, which are time-consuming and resource-intensive [22]. The process of assembling diet matrices is particularly challenging, requiring synthesis of diverse data sources including field studies, literature reviews, and expert opinion. This creates a significant bottleneck in ecosystem model development, especially when applying models to new geographical contexts [23]. Recent advances in artificial intelligence (AI) offer new opportunities to streamline the food web construction process and avoid such bottlenecks [39]. AI tools have demonstrated success in both knowledge/evidence synthesis tasks [40, 26, 4, 38, 45, 33], ecological and environmental tasks [13, 29, 7, 12, 32] and modelling tasks [28, 42, 25], but their application to process-based ecosystem modelling remains nascent. The key challenge lies in ensuring that AI-driven approaches can effectively synthesise available information while maintaining ecological validity.

We present “Synthesising Parameters for Ecosystem modelling with LLMs (SPELL), a novel and flexible framework for assembling and synthesizing user-defined and online resources to construct food webs using Large Language Models (LLMs). Our approach integrates multiple data sources, including global biodiversity databases, species interaction repositories, and locally-held unstructured or structured text, to automate key steps in food web development. SPELL employs user-selected LLMs to group species into functional units and estimate trophic interaction strengths. We evaluate the system in four distinct Australian marine ecosystems the Northern Australia, South East shelf, South East offshore, and Great Australian Bight regions, where we assess the reproducibility of the approach, and in the Great Australian Bight where we assess the accuracy of the approach. Specifically, we test the precision (repeatability) and scientific accuracy of automated species grouping decisions, and the precision and accuracy of the resulting diet matrix proportions, with accuracy defined in terms of similarity to expert estimates. These regions offer contrasting environmental conditions, species assemblages, and ecological dynamics, providing a robust test of SPELL’s adaptability and reliability.

## 2. Methods

### 2.1. SPELL Overview

The development of food web models requires substantial time organizing species into functional groups and determining their trophic interactions. SPELL automates these tasks through a five-stage process that integrates artificial intelligence with ecological databases (Figure 1).

**Figure 1:**
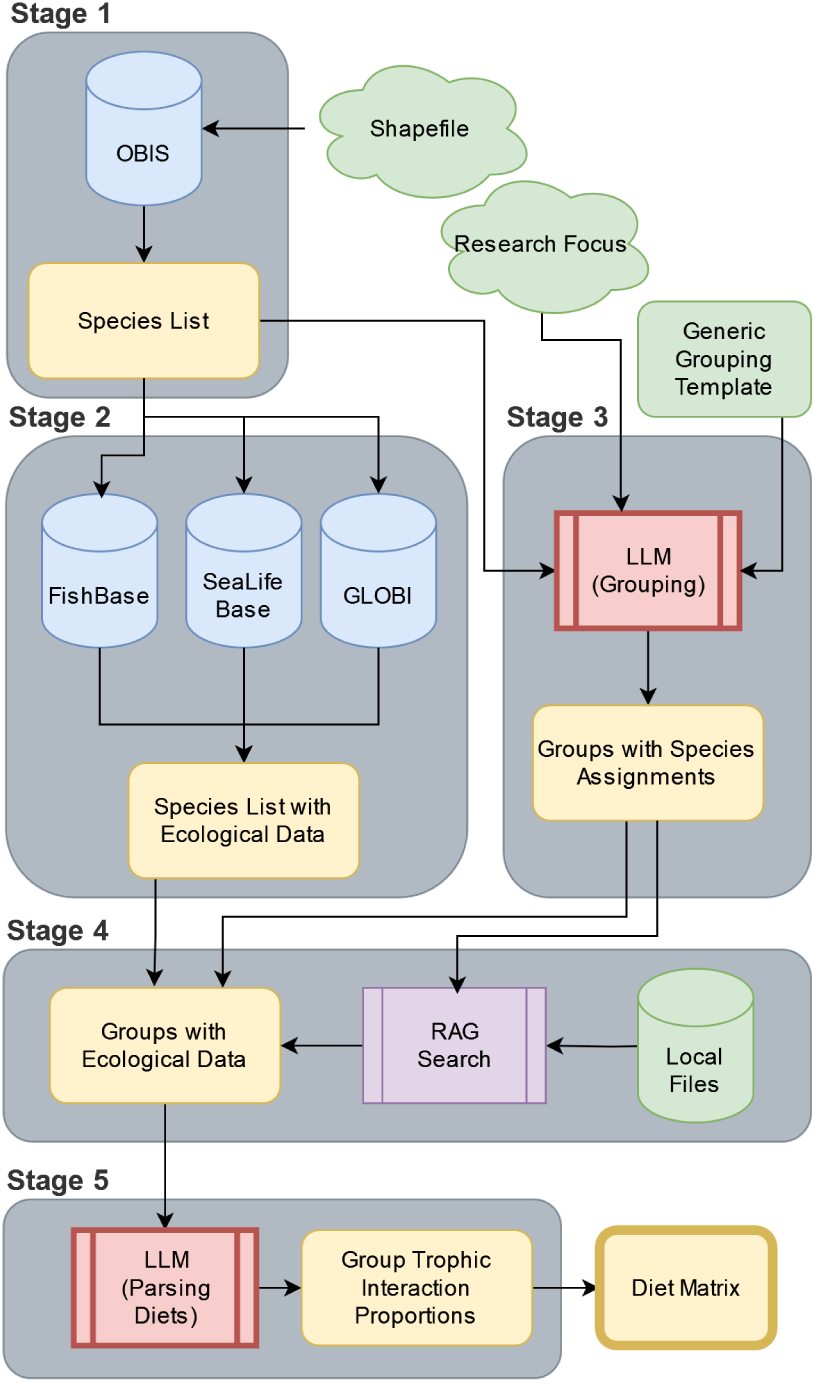
Overview of SPELL for food web construction. The process consists of five stages: species identification, ecological data collection, functional group organization, diet data synthesis, and food web construction. Each stage integrates multiple data sources and analytical approaches, with user-provided inputs (shown in green) guiding key decisions.

The first step in the process is to define a model domain and the resultant shapefile is used to derive a comprehensive species list from Ocean Biodiversity Information System (OBIS) [18]. In Stage 2, this species list is enriched with ecological data from FishBase and SeaLifeBase [14], which provide life history traits, ecological parameters, and diet information, and with trophic interactions from the Global Biotic Interactions (GLOBI) database [34].

Stage 3 employs a LLM for species grouping. We use Claude Sonnet-3.5 (hereafter referred to as ‘Claude’)[1], though other LLMs can be incorporated. The grouping process considers both the research focus specified by the user and a generic grouping template, assigning species into functional groups.

Stage 4 uses a retrieval augmented generation (RAG) system (e.g. [26]) to synthesize the species-level ecological data into group-level summaries and incorporates user-provided local knowledge from a vector storage database (a database that stores and retrieves information based on semantic similarity rather than exact text string matching), Chroma [9], to produce group-level diet composition estimates. Finally, Stage 5 uses the LLM to parse and structure this combined data into a food web matrix, determining trophic interaction proportions among functional groups.

Section S4 of the supplementary material contains detailed documentation of all processing steps, including database queries, literature search criteria, and ecological classification rules. The complete codebase and configuration files can be found at https://github.com/s-spillias/AI-EwE-Diets.

#### 2.1.1. Species Identification

SPELL begins by taking a user-defined shape file that defines the study region boundaries. It then accesses the Ocean Biodiversity Information System [18] through the robis R package [6], which enables automated querying and data processing. We chose OBIS as our primary data source due to its extensive marine species coverage and standardized taxonomic classifications. SPELL uses the ‘checklist’ function from robis to retrieve scientific names and complete taxonomic classifications from kingdom to species level for all recorded species within these boundaries. Whilst collecting all occurrences would help with estimating distributions and biomasses for other modelling purposes, the time and computational resources required to process such large datasets are prohibitive and so we focus on presence only.

To limit the amount of processing required by the LLM, the raw OBIS data is transformed in two steps. First, the R script filters the dataset using OBIS’s is_marine flag to eliminate terrestrial species that may occur in coastal records. Second, it removes taxonomic redundancy using a rank-based approach that retains only the most specific classification level available. For example, if our dataset contains both *Pagrus auratus* (species-level) and *Pagrus* (genus-level) entries for the same organism, our algorithm retains only the species-level entry. Our approach processes taxonomic ranks from most specific (scientific name) to most general (kingdom), keeping only the entry with the highest taxonomic resolution for each organism. Higher-level taxa (e.g., genus, family) are excluded when a species-level entry is available; they are not merged or assumed to represent the same species. If no species-level record exists, the most specific available rank is retained.

The final species list is stored in a structured CSV file containing verified marine species and their complete taxonomic hierarchies. The complete R implementation, including rank-based filtering algorithms and geographic processing functions, is available in the project repository.

#### 2.1.2. Data Harvesting

Following species identification, SPELL gathers ecological and life history information for each identified species. From SeaLifeBase and FishBase [14], it extracts a range of information, including habitat preferences (marine, brackish, or freshwater), depth range distributions, maximum body lengths, and diet data. These databases are accessed through their publicly available PARQUET files using DuckDB for efficient querying of large datasets.

Our species filtering protocol applies constraints to meet Ecopath with Ecosim modelling requirements. We retrieve presence records from OBIS within the study region and remove taxonomic redundancy by retaining only species-level entries (two-word scientific names). Higher-level taxa (genus or family) are excluded to ensure downstream linkage to species-specific database identifiers (but are retained in the OBIS output for record-keeping purposes). For diet composition, we extract food item data from FishBase and SeaLifeBase keyed by species-level SpecCodes, including prey codes, food groups, and prey stage (as is found in FishBase and SeaLifeBase; see https://github.com/s-spillias/AI-EwE-Diets). We specifically filter for juvenile and adult life stages, as larval stages are typically incorporated into planktonic functional groups rather than treated as separate components of adult diets.

We supplement the base biological data with interaction information from the Global Biotic Interactions (GLOBI) database [34]. For each species, we query the GLOBI API using URL-encoded species names to retrieve interaction records in CSV format. The GLOBI data processing preserves the raw interaction data and treats directional relationships (‘eats’/‘preysOn’ and ‘eatenBy’/‘preyedUponBy’) as complementary evidence of trophic interactions. For each predator-prey group pair, we tally the total number of observed interactions, which provides information about the relative frequency of feeding relationships between groups. We further enrich this data through retrieval-augmented generation (RAG) searches of regional literature (detailed in Section S4.2 of the supplementary material), focusing on specific feeding relationships and dietary preferences.

Technical implementation details are provided in Section S1 of the supplementary material.

#### 2.1.3. Species Grouping

We implemented a template-based approach where the LLM is provided with an *a priori* list of possible, generic functional groups (template) and is instructed to assign taxonomic groups to those groups with permission to expand the list if it cannot match a taxonomic group to one of the provided functional groups, and the possibility of returning fewer groups than are provided. By default, SPELL uses a user-defined grouping template (provided in Section S1 of the supplementary material) that leverages a user’s ecosystem modelling experience while allowing for regional customization by either the user or LLM. Due to the complexity of defining ecological groups for EwE models, we have implemented additional templategeneration options but do not use them for validation in this study (See the https://github.com/s-spillias/AI-EwE-Diets for more details). These functional groups are broad ecological units designed for ecosystem modeling and are not strict trophic guilds. While diet is one factor considered, grouping also accounts for habitat, size, and other ecological traits to capture overall functional roles.

Because OBIS can return thousands of species for a given region, instead of using an LLM to classify each species individually, which is timeand cost-prohibitive, we group species hierarchically to reduce the number of classifications required. SPELL does this iteratively by traversing the resulting OBIS database, from kingdom to species, classifying taxonomic groups into functional groups at finer and finer resolutions. Starting at the Kingdom level, the LLM is asked to classify taxa into functional groups. Taxa that the LLM does not think fall neatly into a specific functional group undergo evaluation at finer taxonomic levels until reaching a definitive group assignment or finally reaching the taxonomic level of species, at which point an assignment must be made.

For example, when classifying something like the Western Australian Dhufish (*Glaucosoma hebraicum*), after passing through the Kingdom Animalia, the phylum Chordata is evaluated. Since Chordata includes diverse feeding strategies from filter-feeding tunicates to predatory fish, the LLM, possessing this knowledge innately from its training, marks it for resolution at a finer level. At the class level, the LLM evaluates Actinopterygii, which is again marked for resolution due to its diverse feeding strategies. Continuing through the taxonomic hierarchy, the family Glaucosomatidae is eventually reached, where all members share similar ecological roles as demersal predators, allowing classification into the demersal carnivore functional group. This hierarchical approach substantially reduces the number of required classifications, although is vulnerable to misclassifications at higher taxonomic levels if the LLM does not have sufficient ecological capability. The success of this approach is highly dependent on the ability of the groups in the template to properly capture the overall ecological relations that are needed to model the research question. Success is also dependent on the LLM’s ability to understand the ecological roles of taxa and is a key target for validation in this study. We provide an initial evaluation of the quality of this LLM-generated grouping in Section 2.2.5.

At each taxonomic level, the LLM evaluates taxa against the selected grouping template using the following prompt (where square brackets indicate dynamically updated variables):

You are classifying marine organisms into functional groups for an Ecopath with Ecosim (EwE) model. Functional groups can be individual species or groups of species that perform a similar function in the ecosystem, i.e. have approximately the same growth rates, consumption rates, diets, habitats, and predators. They should be based on species that occupy similar niches, rather than of similar taxonomic groups.

Examine these taxa at the [rank] level and assign each to an ecological functional group.

Rules for assignment:

- If a taxon contains members with different feeding strategies or trophic levels, assign it to ‘RESOLVE’
- Examples requiring ‘RESOLVE’:

– A phylum containing both filter feeders and predators
– An order with both herbivores and carnivores
– A class with species across multiple trophic levels

- If all members of a taxon share similar ecological roles, assign to an appropriate group
- Only consider the adult phase of the organisms, larvae and juveniles will be organized separately
- Only assign a definite group if you are confident ALL members of that taxon belong to that group

Taxa to classify: [List of taxa]

Available ecological groups (name: description): [List of available groups and their descriptions]

Return only a JSON object with taxa as keys and assigned groups as values.

When the research focus indicates groups requiring higher resolution (e.g., commercial fisheries species, or a specific species of conservation concern), the following additional guidance is added to the prompt:

SPELL maintains complete provenance information, including the source of group definitions and any AI-suggested modifications. SPELL automatically includes a Detritus functional group to represent non-living organic matter in the ecosystem. For fisheries-related work, users could also include a discards group that is split off of this general’Detritus’ category. Finally, a detailed grouping report is produced which documents all of the classification decisions for later human review. This allows for a human user to quickly assess the LLM’s decision-making and flag any potential mistakes.

#### 2.1.4. Food Web Construction

After species are grouped into functional groups using SPELL, we reassign their diet and ecological data, sourced from FishBase, SeaLifeBase, and GLOBI, to these new groups. Because each source describes diets differently, we developed a standardized approach to make the information compatible and comparable across all groups.

GLOBI provides species-level predator-prey interactions, which we translate into functional group-level relationships. We start by checking whether each prey species already has a known group assignment. If not, we examine the species’ taxonomic hierarchy, moving from species to genus, family, and higher ranks, to find the closest match among our existing group assignments. When taxonomic information is incomplete or ambiguous, we apply namecleaning rules to assign the prey to the most plausible group. This process ensures that all GLOBI interactions are consistently expressed in terms of our functional groups.

In contrast, FishBase and SeaLifeBase describe diets using hierarchical food categories, such as “zooplankton > copepods” or structured fields like FoodI, FoodII, and FoodIII. We simplify these by collapsing the hierarchy into the most specific and informative category available, while removing vague or placeholder terms like “N.A.” or “others.” These cleaned categories are then matched to the same set of functional groups used for the GLOBI data.

Once all sources are standardized, we compile a comprehensive diet profile for each functional group. This profile integrates prey group counts from GLOBI, cleaned food categories from FishBase and SeaLifeBase, and textual diet descriptions retrieved through retrieval-augmented generation (RAG). We then use the LLM to synthesize this information into a structured summary of diet composition for each group. We also pass a list of example species from a group by compiling one species from each of the three most common genera within a given group. The following prompt guides this first LLM analysis:

Sometimes these responses contain functional groups that are not included in the list of accepted groups or do not add up to 100%. Therefore, the initial diet summaries are passed to a second LLM step that standardizes the proportions and maps any yet undefined group to the already-defined functional groups. This second step converts the approximate summaries into a structured food web matrix, with prey items as rows and predators as columns. Each cell contains the proportion of the predator’s diet comprised of that prey item. The food web matrix is then output as a CSV file for use in ecosystem models and food web analyses.

When prey items do not exactly match functional group names, we employ a hierarchical matching system. SPELL first attempts exact matches, then falls back to case-insensitive partial matching using species names. For example, if the AI returns a prey item “snapper” that doesn’t exactly match any functional group, SPELL would match it to a functional group containing “snapper” in its name such as “Pink Snapper”.

This process is the second target of validation in this study and is evaluated in Section S2. The complete codebase and configuration files are available at https://github.com/s-spillias/AI-EwE-Diets.

### 2.2. Validation

Our validation framework assesses both the precision (consistency across multiple runs for the same food web) and accuracy (comparison with expert-created food web matrices) of the LLM-driven model construction process. We used four distinct Australian marine regions for these assessments: all four regions (Northern Australia, South East shelf, South East Offshore, and Great Australian Bight) to evaluate precision, with the Great Australian Bight also used to evaluate accuracy.

We executed model generation across three distinct phases. In phase one, we established baseline configurations for each study region by processing species occurrence data and downloading relevant species data. In phase two, where the LLM is first called, we executed five independent iterations per region, maintaining fixed input parameters while allowing the LLM’s stochastic decision processes to generate variation in outputs for grouping assignment and diet proportion synthesis. For all of these validation runs, the resulting functional groups are the product of the two-step process described in Section 2.1.3 where we provide a generic candidate template of functional groups and their descriptions (developed by the co-authors for this study; Table S1), then the LLM assigns species found in phase one to these generic groups, with the flexibility to modify or create new groups when taxa do not fit existing categories. In phase three, we conducted detailed statistical analyses of both precision across iterations and accuracy compared to expertcreated matrices.

For the precision assessment, we calculated Jaccard similarity coefficients between all possible pairs of iterations to evaluate the consistency of species groupings and diet matrices. We also assessed the stability of diet proportions using a normalized deviation-based stability score, and evaluated prey composition consistency using Spearman’s rank correlation. We applied KruskalWallis tests to region-level summaries of diet stability and prey composition correlations to test for significant differences in trophic structure across ecosystems. Additionally, to confirm that the LLM-generated groupings were not random, we conducted a chi-square goodness-of-fit test comparing the aggregated distribution of species across functional groups to a uniform distribution. For the accuracy assessment, we averaged the diet proportions of the five LLM-generated matrices and compared the resulting matrix with an expert-created matrix for the Great Australian Bight ecosystem (C. Bulman pers. comm.) that was used to inform [15].

### 2.2.1. Study Regions

We selected four Australian marine regions that present distinct ecological characteristics and modelling challenges for SPELL (Figure 2): the Northern Australia region, which represents a tropical ecosystem characterised by a broad shelf and complex mix of reef systems, seagrass meadows, mangrove forests and bare sediment communities with seasonal monsoon influences; the South East shelf region, a temperate coastal system with a network of rocky reefs and kelp forests, rapidly changing environmental conditions due to climate change, comprehensive diet information in established databases, well-documented EwE models spanning multiple years, and active research programs; and the South East Offshore region, a deep-water ecosystem that challenges SPELL with data-limited conditions and unique ecological patterns relating to oceanic through flow, shifting current patterns, low productivity patches interspersed with production concentrating canyons and seamounts. For these three systems we provided a generic research focus: “Future of Seafood”, to get a sense of SPELL’s general behaviour.

**Figure 2:**
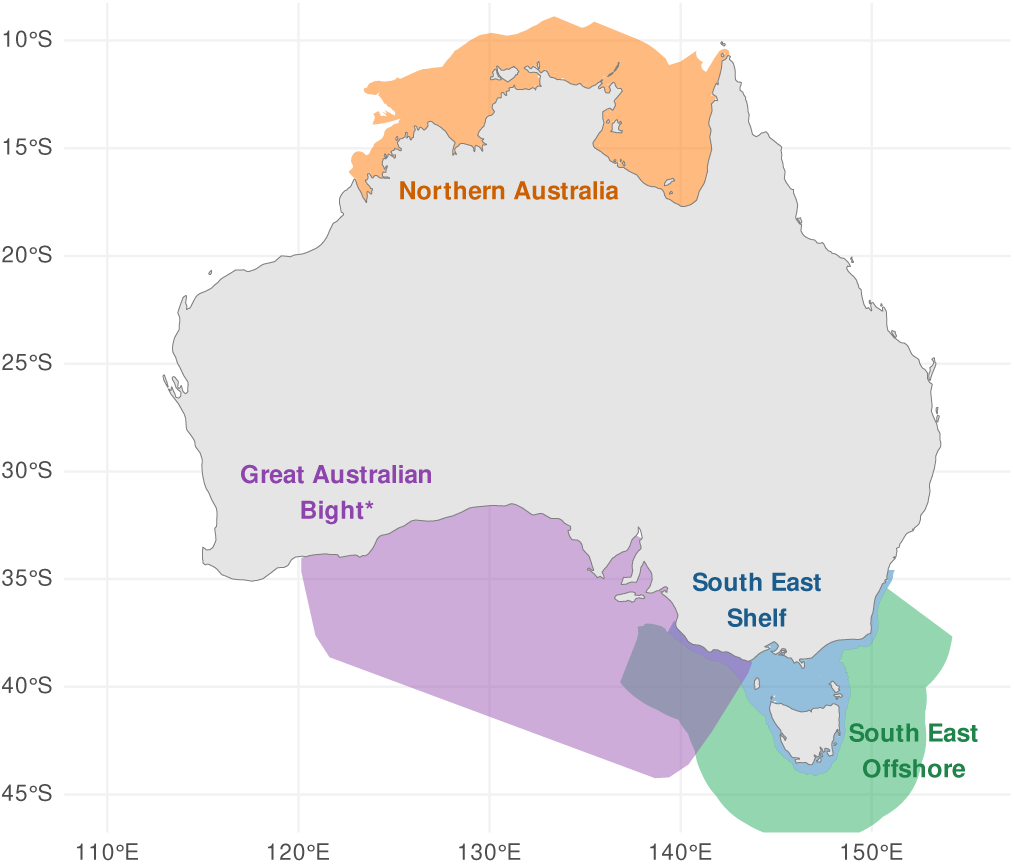
Map of the four study regions used to validate our approach: three of the regions were used to test the precision (consistency) of the approach: Northern Australia (orange), South East shelf (blue), South East Offshore (green). We used the fourth region, the Great Australian Bight (purple) to test the accuracy of our approach by comparing it to the initial food web matrix used in [15]

For accuracy assessment, we used the fourth region: the Great Australian Bight (GAB), which represents a region of high conservation significance spanning diverse habitats. The GAB has been extensively studied, with research characterising the shelf and slope ecosystems from phytoplankton through to marine mammals and birds [17, 15]. We used this region to compare the resulting diet matrix from our process to the expert-created matrix used in [15], allowing us to evaluate the accuracy of our approach. For this assessment, we used a more specific research focus, derived from the abstract of [15]:

The ecological consequences (changes in trophic linkages and biomass flow) of increased fishing pressure, shipping, ocean warming, spatial reserves and oil spills due to ship collision.

The contrasting characteristics of these four regions provide a robust test of SPELL’s adaptability across different ecological contexts.

#### 2.2.2. Grouping Precision Analysis

To assess the precision of SPELL-generated species groupings, we developed quantitative measures of grouping precision. For each of the four regions, we conducted five independent iterations, resulting in five grouping outcomes per region. We analyzed precision within each region separately.

For each region, we tracked each species’ group assignments across the five iterations and calculated a precision score:

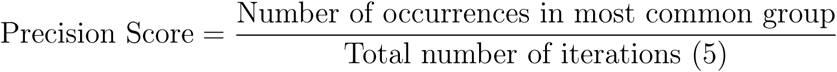

This metric quantifies SPELL’s decision-making reliability for individual species within a specific ecological context. We classified species with precision scores below 1.0 as unstable, indicating variable group assignments across iterations. A species with a precision score of 1.0 was assigned to the same functional group in every iteration, demonstrating high precision in SPELL’s decision-making.

While the precision score measures stability at the individual species level, we also needed to evaluate stability at the group level. To do this, we assessed group stability using the Jaccard similarity coefficient between all possible pairs of iterations within each region:

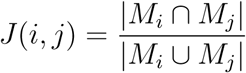

where *M_i_* and *M_j_* represent the sets of species members in iterations *i* and *j*. Unlike the precision score, which focuses on whether individual species are consistently assigned to the same group, the Jaccard similarity measures whether a group consistently contains the same set of species across iterations. For example, a group might maintain a stable core of species while experiencing minor variations in peripheral members, which would be captured by the Jaccard similarity but not by individual species precision scores.

For each region, we calculated the overall stability score by averaging Jaccard similarities across all possible pairs of iterations. This approach reveals how consistently SPELL identifies and maintains ecologically meaningful groupings across different runs. We also conducted a chi-square goodnessof-fit test comparing the aggregated distribution of species across functional groups to a uniform distribution, which evaluates whether certain groups consistently receive more species than others, indicating structured ecological reasoning rather than random allocation.

#### 2.2.3. Grouping Accuracy Assessment

To assess the ecological validity of SPELL-assigned functional groups, we conducted a manual validation of taxonomic grouping decisions. We did not use the original expert groupings from [15] for two reasons: first, SPELL handled many more taxonomic groupings and species than the human-created groupings, and second, comprehensive data for each functional group was no longer available to do a direct comparison. Therefore, we undertook a direct evaluation where the lead author, SS, reviewed each taxonomic entity (ranging from species to higher taxonomic levels) for a single iteration of the GAB (n=675 SPELL-decisions) and evaluated whether SPELL had correctly assigned it to an appropriate functional group based on known ecological characteristics.

For each taxonomic group assigned by SPELL to a functional group, we researched the known ecological characteristics of that taxon, including feeding behavior, habitat preferences, and trophic position. We compared these ecological characteristics to the description of the functional group provided in the grouping template. Based on this comparison, we classified each assignment as either “Correct” (the taxon fits well within the functional group), “Partial” (the taxon partially fits the functional group but has some characteristics that don’t align), “Incorrect” (the taxon was inappropriately assigned), or “Unsure” (insufficient information was available to make a determination).

#### 2.2.4. Diet Matrix Precision Assessment

To evaluate the precision of SPELL-generated trophic interactions and assess SPELL’s ability to capture distinct ecological patterns, we developed a multi-metric analysis approach. For each region separately (with five iterations per region), we calculated the following metrics for each predator-prey interaction:

1. Presence ratio across iterations:

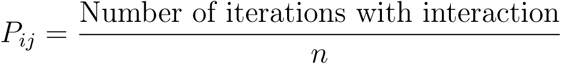

where *n* is the total number of iterations (5 per region), and an interaction is present when the diet proportion *x_ijk_ >* 0 for predator *i* consuming prey *j* in iteration *k*.

2. Mean diet proportion:

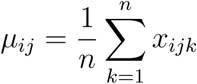

where *x_ijk_*represents diet proportion for predator *i* consuming prey *j* in iteration *k*.

3. Stability score:

We first calculate a normalized deviation score for each predator-prey interaction:

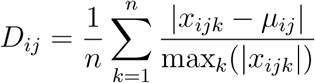

where *µ_ij_* is the mean diet proportion across iterations, and max*_k_*(|*x_ijk_*|) is the maximum absolute value across iterations. Because diet proportions (*x_ijk_*) are bounded between 0 and 1, this deviation score ranges from 0 to 0.5.

Then, to create a more intuitive stability score where higher values represent greater stability, we invert this measure:

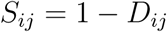

This transformation yields a stability score bounded between 0.5 (maximum instability) and 1 (perfect stability), with higher values indicating more consistent diet proportions across iterations.

We chose this stability metric over traditional variance measures for several reasons. First, by normalizing deviations by the maximum value, the metric achieves scale independence, allowing meaningful comparisons between interactions of different magnitudes. For example, the sequences [0.2, 0.2, 0.2, 0.2, 0.1] and [0.02, 0.02, 0.02, 0.02, 0.01] would yield the same stability score despite having different absolute variances. Second, the bounded range between 0 and 1 provides an intuitive scale for assessing stability, unlike the unbounded nature of variance. Third, when diet proportions are of similar magnitude across iterations, this approach prevents minor fluctuations in small values from disproportionately influencing the stability assessment. However, in cases where iterations contain both very small and substantially larger values, the scale independence property means the stability assessment will be more sensitive to relative deviations in the larger values.

We classified interactions as unstable when their stability score fell below 0.7, corresponding to a normalized deviation of 0.3 in the original metric. This approach balances sensitivity to meaningful ecological variation while avoiding flagging minor fluctuations that are expected in complex ecological systems. Variations in predator-prey interaction strengths beyond this threshold suggest fundamental uncertainty in the trophic relationship that would propagate through ecosystem simulations and affect model predictions. To illustrate this metric:

- A stable interaction (S = 0.92) might show values [0.02, 0.02, 0.02, 0.02, 0.01], where proportions remain very similar across iterations
- An unstable interaction (S = 0.61) might show values [0.027, 0.25, 0.25, 0.067, 0.25], where proportions vary substantially between iterations, indicating inconsistent characterization of the predator-prey relationship by roughly an order of magnitude

This metric provides a continuous measure of stability that handles both presence/absence patterns and magnitude variations in a unified way. To assess SPELL’s ability to capture distinct ecological patterns across regions, we employed pairwise Spearman correlations between iterations to evaluate the precision of predator-prey relationships. This non-parametric approach accounts for the potentially non-normal distribution of diet proportions. We supplemented this with Kruskal-Wallis tests to identify significant differences in trophic structure across regions, providing evidence of SPELL’s ability to distinguish unique ecological characteristics in different marine ecosystems.

To assess SPELL’s ability to capture distinct ecological patterns across regions, we also evaluated composition-level reliability using Spearman’s rank correlation (*ρ*) for each predator across all pairs of iterations 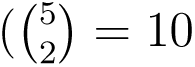 pairs per predator). This non-parametric measure captures the stability of prey rank ordering within predator diets, independent of absolute diet proportions. We first aligned all matrices by common predator rows and prey columns across iterations and row-normalized each matrix to proportions so that comparisons reflect relative composition rather than absolute biomass. For each predator, we calculated *ρ* between its prey composition vectors for every iteration pair. Comparisons with fewer than two valid prey entries or constant vectors (no rank variation) were excluded, as Spearman’s *ρ* is undefined in these cases.

To summarize reliability at the region level, we applied Fisher’s *z*-transformation to all valid *ρ* values, computed the mean *z*, and back-transformed to obtain the mean correlation. We estimated uncertainty using non-parametric bootstrap resampling (2,000 replicates) to derive 95% confidence intervals.

#### 2.2.5. Diet Matrix Accuracy Assessment

To evaluate the accuracy of SPELL-generated diet matrices against expert knowledge, we conducted a detailed comparison using the Great Australian Bight (GAB) ecosystem model developed by Fulton et al. [15]. We obtained the original, unbalanced diet matrix constructed by expert marine ecologists (C. Bulman, personal communication) and compared it with five independently generated SPELL matrices for the same region. To specifically assess the diet proportion accuracy, we provide SPELL with a grouping template consisting of the same list of groupings from the GAB model, thus testing SPELL’s ability to sort species into the correct groups and then assign diet proportions according to those groups.

The analysis examined two fundamental aspects of the diet matrices: the structural patterns of predator-prey relationships and the quantitative diet proportions. For the structural (presence-absence) comparison, we flattened the 59 by 59 predator-prey adjacency matrices to binary vectors after restricting to the set of functional groups common to both matrices (59 groups; 3481 potential links). We defined a link as present when the mean diet proportion (averaged across five SPELL iterations) was > 0. We quantified overall agreement and Cohen’s Kappa (*κ*), which corrects for chance agreement and is appropriate for categorical outcomes. We also report the observed agreement proportion (Po), defined as:

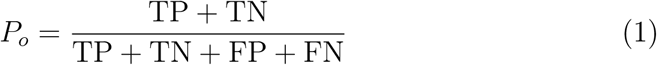

where TP and TN are the counts of true positives and true negatives, and FP and FN are false positives and false negatives, respectively.

For predator-prey pairs where both matrices indicated an interaction, we conducted quantitative comparisons of the diet proportions. We compared diet proportions only for predator-prey pairs where both matrices indicated a link (common links). We report Pearson correlation coefficients (*ρ*) and the distribution of absolute differences in proportions (mean, median, quartiles). We chose this measure because, unlike the stability metrics used in the precision assessment which evaluate consistency across multiple iterations, correlation analysis is specifically designed to quantify the alignment between two distinct matrices the SPELL-generated and expert-created matrices. This approach directly addresses the accuracy objective by measuring how well the SPELL-generated diet proportions correspond to expert knowledge, rather than measuring consistency across multiple SPELL-generated iterations. We performed these analyses both at the whole-matrix level and for individual predator groups, enabling identification of systematic patterns in SPELL’s performance across different taxonomic groups.

## 3. Results

Our validation framework assessed three key aspects of the AI-assisted ecosystem modelling approach: reproducibility of species groupings, consistency of food web construction, and accuracy against expert-derived matrices.

### 3.1. Species Grouping Precision

#### 3.1.1. Classification Consistency Analysis

SPELL successfully reduced ecological complexity while preserving meaningful biological relationships. Starting with 63 potential functional groups provided in the default template (See S4), it identified 34-37 region-specific groups. Chi-square tests confirmed the non-random nature of these groupings, showing consistent species assignments across all regions (p < 0.001). This statistical significance provides evidence that SPELL makes systematic grouping decisions rather than arbitrary assignments.

SPELL achieved high classification stability for groups across all regions. Mean consistency scores, where 1.0 represents identical species assignments to groups across all groups and within-region iterations, were exceptionally high: 0.997 for both Northern Australia and South East shelf, and 0.998 for South East Offshore. This translated to very low proportions of species that were variably classified across the five iterations: only 0.99% (103 species) in Northern Australia, 1.06% (125 species) in South East shelf, and 0.73% (87 species) in South East Offshore. These results demonstrate that SPELL’s classifications remained stable despite the stochastic nature of the AI decision-making process.

Among the small percentage of variably classified species, we identified consistent patterns of classification instability (Table 1). These species typically oscillated between ecologically similar functional groups, such as macrozoobenthos and benthic infaunal carnivores in the Northern Australia, or piscivores and deep demersal fish in the South East Shelf region. This suggests that classification uncertainty occurs primarily at ecological boundaries where functional roles overlap.

**Table 1:** Dominant patterns of species classification instability across three study regions. The table presents the most frequent oscillation patterns between functional groups for species that were inconsistently classified across the five framework iterations. For each region, the total number of variably classified species is shown (representing less than 1.1% of all species), along with the percentage distribution of different oscillation patterns.

| Region | Most Common Pattern | Count | % of Total |
| --- | --- | --- | --- |
| Northern Australia<br>(103 species) | Macrozoobenthos ↔ Benthic infaunal carnivores | 28 | 27.2% |
| | Benthic filter feeders ↔ Deposit feeders | 25 | 24.3% |
| | Prawns ↔ Macrozoobenthos | 21 | 20.4% |
|  | Other patterns | 29 | 28.1% |
| South East Shelf<br>(125 species) | Piscivores ↔ Deep demersal fish | 42 | 33.6% |
| | Benthic grazers ↔ Benthic carnivores | 31 | 24.8% |
| | Planktivores ↔ Mesopelagic fish | 28 | 22.4% |
|  | Other patterns | 24 | 19.2% |
| South East Offshore<br>(87 species) | Benthic filter feeders ↔ Benthic carnivores | 25 | 28.7% |
| | Macrozoobenthos ↔ Deep demersal fish | 22 | 25.3% |
| | Mesozooplankton ↔ Macrozoobenthos | 18 | 20.7% |
|  | Other patterns | 22 | 25.3% |
| GAB<br>(87 species) | Benthic infaunal carnivores ↔ Macrozoobenthos | 32 | 36.8% |
| | Benthic filter feeders ↔ Deposit feeders | 28 | 32.2% |
| | Prawns ↔ Benthic infaunal carnivores | 15 | 17.2% |
|  | Other patterns | 12 | 13.8% |

*Note:* Arrows indicate group assignment oscillation between iterations. Complete species-level data available in Section S3 of the supplementary material.

**Table 2:** Computational requirements by region and processing stage.

| Region | Species<br>Count | Processing Time (hours) |  |  |  |  |
| --- | --- | --- | --- | --- | --- | --- |
|  |  | Identification | Data<br>Download | Grouping | Diet<br>Collection | Matrix<br>Construction |
| Northern Australia | 10,621 | 0.01 | 2.2 | 0.2 | 0.2 | 0.04 |
| South East Shelf | 17,068 | 0.01 | 2.8 | 0.2 | 1.6 | 0.04 |
| South East Offshore | 12,033 | 0.01 | 3.3 | 0.4 | 0.3 | 0.04 |
| Great Australian Bight | 6,957 | 0.01 | 1.3 | 0.3 | 0.5 | 0.07 |

The Jaccard similarity indices reveal high overall stability in group membership across all four regions (Figure 3), with most functional groups showing indices above 0.95. The groups labelled in the figure represent those groups with lower stability indices.

**Figure 3:**
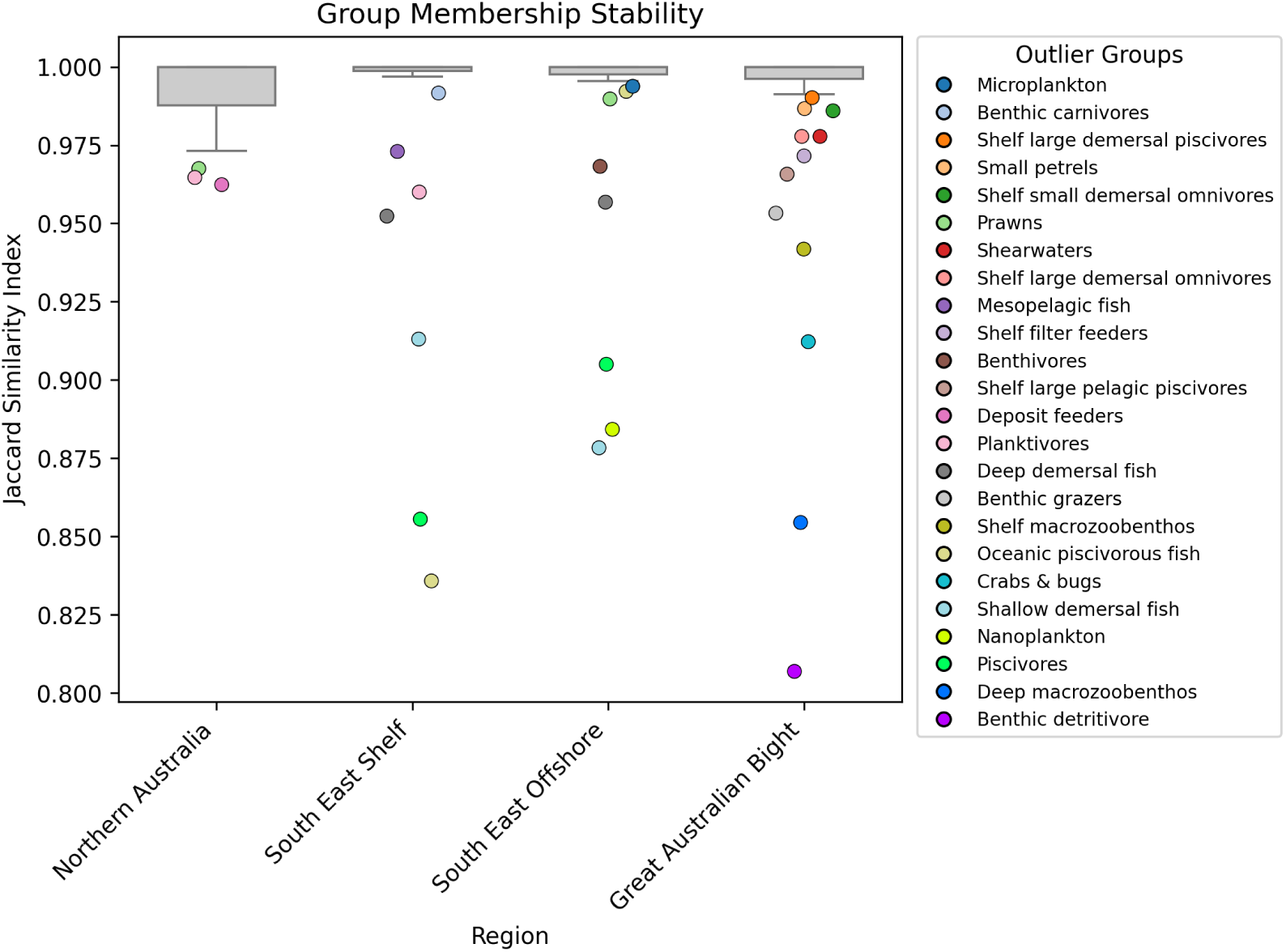
Group membership stability across four regions measured by Jaccard similarity index (0.85-1.0). Most groups show high stability (>0.95), with labelled points out-of-distribution outliers that exhibit lower stability.

Further detailed analysis of group stability patterns across regions is provided in S3.

### 3.2. Food Web Precision

#### 3.2.1. Trophic Interaction Consistency

SPELL identified consistent trophic relationships across all regions, with Northern Australia showing 358 interactions (58.4% having stability scores > 0.7), South East shelf 380 interactions (51.3% of which were stable), and South East Offshore 477 interactions (56.0% stable). As shown in Figure 4, the distribution of stability scores across regions demonstrates that most interactions cluster above the 0.7 threshold, with 117 interactions across regions achieving near-perfect stability (scores >= 0.95). Kruskal-Wallis tests indicated no significant differences in diet stability across regions (p > 0.05), suggesting consistent trophic structure precision. Spearman rank correlations of prey composition between iterations indicated strong reliability across all regions, though with some variation in magnitude. Northern Australia exhibited the highest consistency (mean *ρ* = 0.945, 95% CI = 0.924-0.962), followed by South East Offshore (*ρ* = 0.887, 95% CI = 0.852-0.913), Great Australian Bight (*ρ* = 0.882, 95% CI = 0.849-0.909), and South East Inshore (*ρ* = 0.867, 95% CI = 0.831-0.898). These results demonstrate that the rank ordering of prey shares within predator diets remains highly consistent across model iterations. Detailed diet matrices for each region are provided in S2.

**Figure 4:**
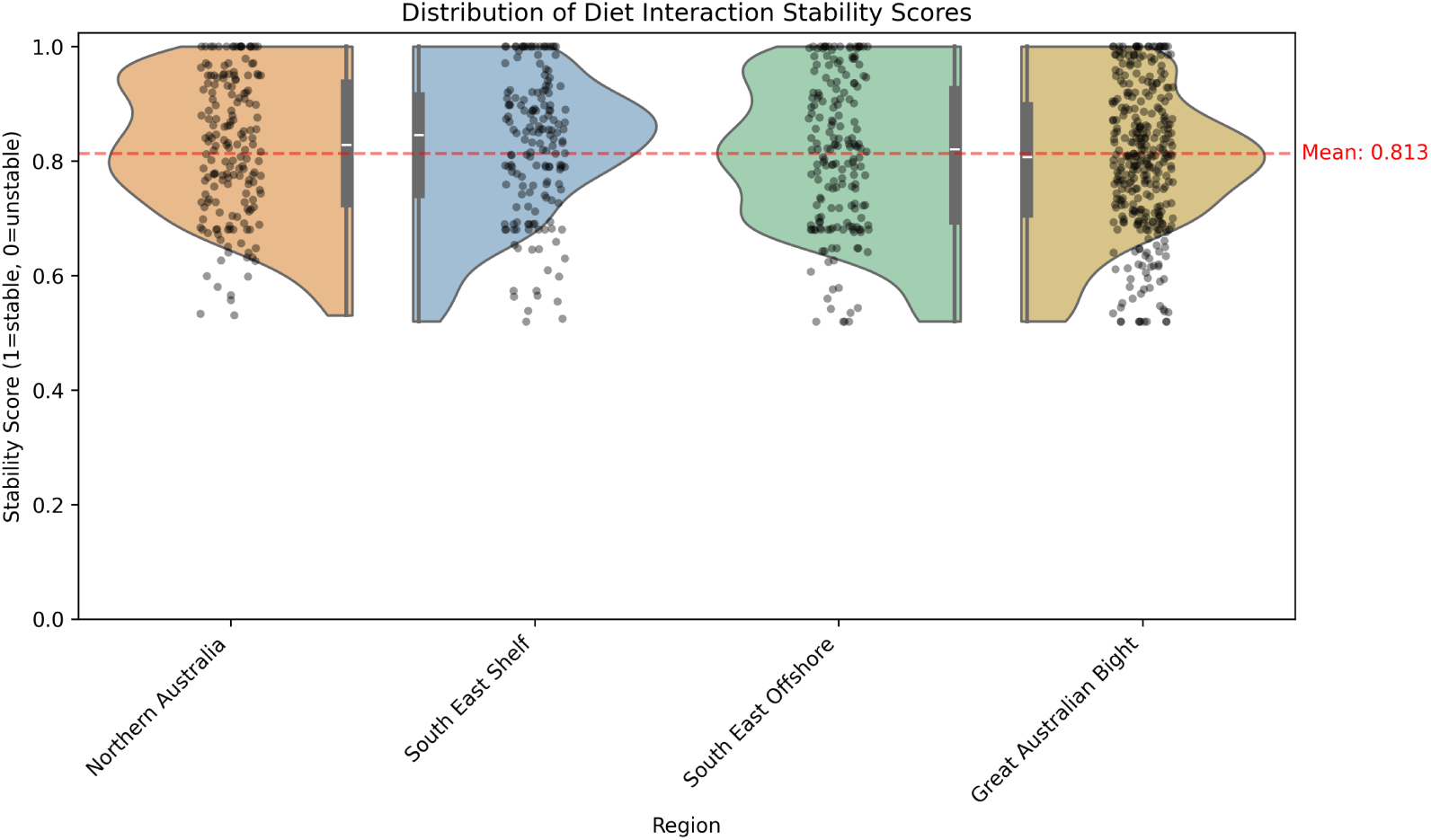
Distribution of diet interaction stability scores across regions for substantial interactions (those comprising more than 5% of a predator’s diet). Half-violin plots show the density of stability scores (1=stable, 0=unstable), with embedded box plots indicating quartiles and median. Individual points represent specific predator-prey interactions, and the red dashed line shows the mean stability score across all regions. The distributions are bounded at one, reflecting perfect stability, with most interactions showing scores above 0.7. Stability scores quantify the consistency of predator-prey interactions across iterations, where a score of 1.0 indicates the interaction was identified with identical diet proportions in all iterations, while lower scores reflect either variable diet proportions or inconsistent identification of the interaction.

### 3.3. Grouping and Diet Proportion Accuracy Assessment: Great Australian Bight Case Study

#### 3.3.1. Taxonomic Grouping Accuracy

To evaluate the ecological validity of AI-assigned functional groups, we conducted a detailed manual validation of 675 taxonomic grouping decisions. The results revealed that 75.3% (508) of the AI’s taxonomic assignments were fully correct, aligning with known ecological characteristics of the taxa. Additionally, 17.3% (117) of assignments were partially correct, where the taxon fit some but not all aspects of the functional group description. For example, these included cases where the AI system designated a taxonomic group as ‘deep’ or ‘slope’ when they might inhabit both, or might designate a taxonomic group as ‘large’ or ‘small’ when members could be one or the other. Only 3.4% (23) of assignments were clearly incorrect (demonstrable incorrect feeding strategy habitats), and 4.0% (27) could not be definitively assessed due to limited ecological information about the taxa (mostly poorly researched deep-water taxonomies).

The accuracy of assignments varied considerably across functional groups (Figure 5). Many functional groups showed perfect or near-perfect assignment accuracy, including all assignments for albatross, pelagic sharks, small phytoplankton, mesozooplankton, small petrels, and several other well-defined groups. These groups typically have clear ecological niches and distinctive characteristics that facilitate accurate classification.

**Figure 5:**
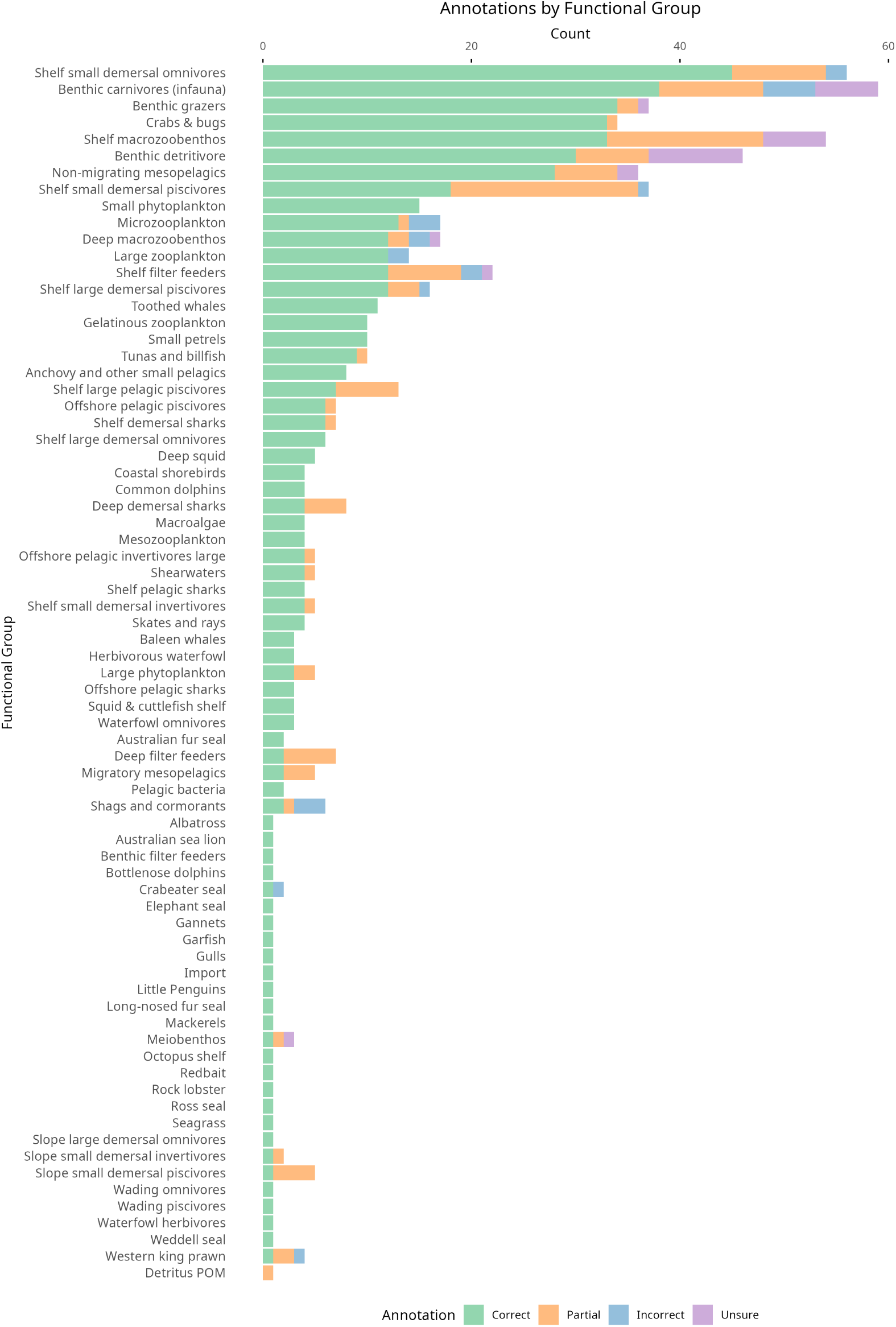
Accuracy of functional group assignments generated by the LLM for the Great Australian Bight case study. The LLM used the template in Table S1 as guidance but could omit unused groups or create new ones. Bars show the proportion of assignments classified as correct (green), partial (orange), incorrect (blue), or uncertain (purple) for each final group.

Functional groups with lower accuracy rates appeared to be constrained by limitations in the AI system’s grouping template, particularly for specialized ecological niches. For example, waterfowl were frequently misclassified into the “Shags and cormorants” group, achieving only 33.3% correct assignments with 50% incorrect assignments. These errors revealed confusion between taxonomically related but ecologically distinct bird groups. Similarly, “Deep filter feeders” showed only 28.6% correct assignments with 71.4% partial assignments, highlighting challenges in classifying deep-sea organisms with complex or variable feeding strategies.

The analysis of partial assignments revealed several recurring patterns. Taxa associated with deep-sea environments (17 taxa) were frequently misclassified, likely due to limited ecological information and the complex nature of deep-sea ecosystems. Parasitic organisms (7 taxa) were also challenging to classify correctly, as they often have complex life cycles that span multiple functional roles. Filter feeders, detritivores, and grazers showed similar patterns of partial classification, typically due to their variable feeding strategies that may change based on environmental conditions or life stage.

#### 3.3.2. Food Web Accuracy

To evaluate SPELL’s accuracy against expert knowledge, we compared its output to an expert-derived Ecopath model of the Great Australian Bight ecosystem [15]. Across the 59 functional groups common to both matrices, SPELL achieved strong overall agreement in the structural (presenceabsence) pattern of trophic interactions (Po = 0.816), with 73.1% of entries correctly identified as absent in both matrices and 8.5% as present in both (TP = 296, TN = 2545, FP = 505, FN = 135). Cohen’s Kappa for the presence-absence comparison was *κ* = 0.381, reflecting moderate agreement once chance agreement is accounted for, consistent with the high class imbalance (positives 12.4%). Despite the prompt’s guidance to retain species of particular interest as individual functional groups, SPELL omitted 17 groups present in the expert matrix, including several commercially important species (Southern Bluefin Tuna, Snapper, King George whiting, and Abalone) as well as Nanozooplankton. Conversely, it generated only two groups not present in the expert matrix (Offshore pelagic invertivores large and Slope large demersal omnivores). This limitation was most evident in commercially important species that typically receive individual attention in expert-created models but were subsumed into broader functional groups by SPELL. A comprehensive visualization of these differences across all functional groups is provided in S2.

As shown in Figure 6a, SPELL demonstrated varying levels of agreement across different functional groups in identifying trophic interactions. The analysis revealed that 8.5% of interactions were present in both matrices (dark purple), while 73.1% were correctly identified as absent in both (light grey). SPELL uniquely identified 14.5% of interactions (teal) that were not present in the expert matrix, while missing 3.9% of expert-identified interactions (yellow). Overall, SPELL achieved an agreement rate of 81.6% with the expert matrix, with a true positive rate (sensitivity) of 0.687 and a true negative rate (specificity) of 0.834.

**Figure 6:**
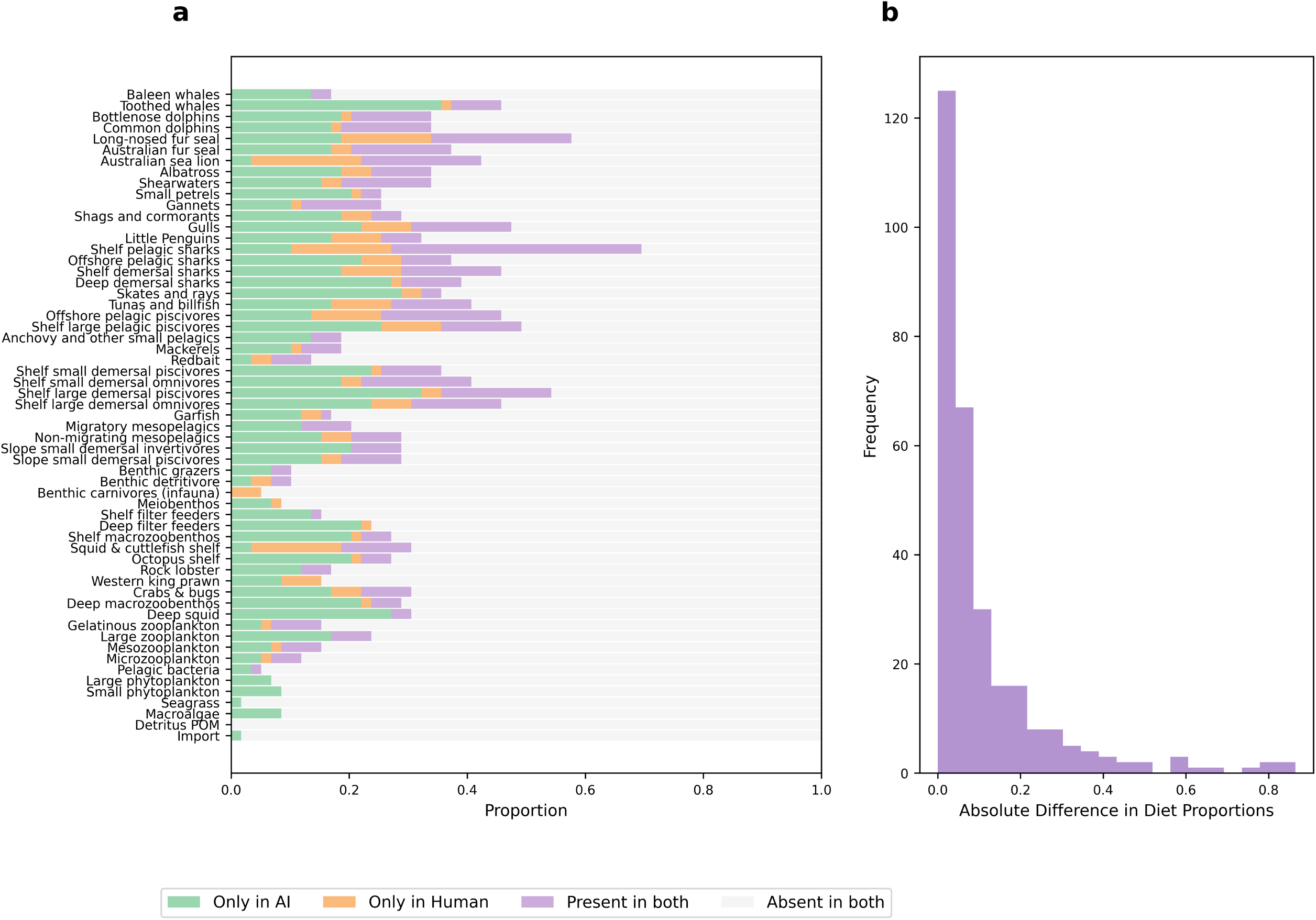
Comparison of expert-created and AI-generated diet matrices for the Great Australian Bight ecosystem. (a) Presence-absence patterns showing the proportion of different interaction types across functional groups. Dark purple indicates interactions present in both matrices, orange shows expert-only interactions, teal shows AI-only interactions, and light grey indicates absence in both. (b) Distribution of absolute differences in diet proportions where both matrices indicate an interaction, showing the frequency of different magnitudes of disagreement between AI and expert estimates.

For interactions present in both matrices (n = 296 common links), quantitative agreement in diet proportions was moderate (Pearson r = 0.385, p < 0.001). Absolute differences were generally small (mean = 0.110; median = 0.058; 25th-75th percentiles = 0.021-0.128), with approximately 80% of differences < 0.2, although a long tail (max = 0.865) indicates notable discrepancies for some predator-prey pairs. This suggests that when SPELL correctly identified a trophic interaction, it often estimated diet proportions within reasonable bounds of expert values, though with notable variations across different predator-prey combinations.

### 3.4. SPELL Implementation and Performance

#### 3.4.1. Scale and Processing Efficiency

We evaluated SPELL through five independent runs across three distinct Australian regions, processing a total of 39,722 species. SPELL handled 10,621 species in the Northern Australia’s tropical reef ecosystem, 17,068 in the South East shelf’s coastal and pelagic environments, and 12,033 in the South East Offshore’s deep-water systems.

#### 3.4.2. Computational Efficiency

Total processing time ranged from 2.2 to 4.8 hours across regions. The most time-intensive stage was the downloading of biological data from online databases, accounting for approximately 69% of the total processing time. Species identification typically required 0.01 hours, while the AIdriven species grouping process averaged 0.26 hours. Diet data collection and matrix construction required 0.7 and 0.04 hours respectively, with final parameter estimation taking 0.20 hours. On average, SPELL required 0.7 seconds per species for data downloading and 0.2 seconds per species for diet data collection, though these rates varied considerably between regions due to differences in data availability and species complexity.

## 4. Discussion

We have shown that SPELL is able to construct food webs for ecosystem models with a fair degree of reliability. This capability addresses a significant challenge in ecosystem modelling, as the increasing use of these models for environmental management and policy decisions requires efficient and accurate development approaches [44, 36]. Constructing reliable ecosystem models traditionally involves complex technical challenges, including species identification, data harvesting, and the creation of accurate trophic interaction networks processes that are particularly demanding in fisheries management contexts where food web models inform ecological and socioeconomic decision-making [5]. SPELL provides a systematic, AI-assisted solution that enhances reproducibility and dramatically reduces the time investment required from months to hours. This efficiency gain aligns with the growing recognition that effective ecological models must balance mechanistic understanding, appropriate spatial and temporal resolution, and uncertainty quantification to support decision-making [36]. By streamlining technical aspects through integration of multiple data sources [8, 11] and AI-driven synthesis [40, 32], our approach allows modelers to dedicate more resources to stakeholder engagement and result communication, potentially increasing the impact of ecosystem modelling on environmental management.

### 4.1. Validation Assessment

Here we have demonstrated SPELL’s potential to generate reproducible, ecologically meaningful components for ecosystem model development while significantly reducing development time. It can complement the traditional approach to model building and expert judgement. SPELL’s ability to construct reliable ecosystem model components reveal both strengths and limitations. SPELL demonstrates strong internal consistency, with high stability scores (99.7-99.8%) in species classifications across regions and robust correlations in predator-prey rankings (*ρ* = 0.87−0.95). This consistency suggests SPELL makes systematic rather than arbitrary decisions in constructing ecological relationships. However, these metrics must be interpreted cautiously, as they reflect SPELL’s reproducibility rather than ecological accuracy.

Our performance metrics are comparable to other recent AI applications in food web modelling, such as FoodWebAI [32], which achieved 90-98% accuracy in correctly determining species’ positions in the food chain (trophic levels) and 74% accuracy in identifying predator-prey relationships (trophic links) across three ecosystems. The grouping accuracy we found was less than others have reported for bespoke chatbots but higher than has been reported for frontier LLMs of the previous generation [35].

The Great Australian Bight comparison against expert knowledge provides important insights into SPELL’s reliability. The high success rate in identifying absent trophic interactions (73.1%) indicates SPELL effectively avoids spurious ecological connections. However, the moderate Kappa coefficient (0.38) with expert-assigned diet proportions and the tendency to miss specialised ecological groups reveals important limitations. SPELL shows a bias toward generalised classifications that may overlook managementrelevant distinctions, particularly when it comes to accommodating functional groups that are comprised of only a single species.

Our detailed taxonomic validation analysis further illuminates SPELL’s ecological accuracy, with 75.3% of taxonomic assignments being fully correct and an additional 17.3% being partially correct. This high overall accuracy rate (92.6% at least partially correct) is consistent with other analyses [32, 12] and suggests SPELL generally makes ecologically sound grouping decisions. However, the analysis also revealed that the success of the grouping may depend on the comprehensiveness of the grouping template that SPELL is provided. Deep-sea organisms, parasitic taxa, and species with complex or variable feeding strategies were more frequently misclassified or only partially correctly assigned. These findings highlight areas where SPELL’s ecological knowledge may be limited or where the predefined functional group templates may not adequately capture or discriminate the full range of ecological roles present in marine ecosystems. Template sensitivity has been examined in a separate benchmarking study [37], which shows how template granularity and batch size influence classification accuracy and offers preliminary guidance on template design for ecological applications. These results also highlight where human triage will be needed when using these methods, showing the kinds of groups where the modeller needs to focus checking or redefine model structures to reduce erroneous assignments. Notably, most classification uncertainty did not stem from clear errors but from taxa that occupy ecological grey zones species whose traits span multiple functional roles. In such cases, SPELL’s variable classifications reflect genuine ecological ambiguity rather than systematic misclassification. This behaviour aligns with principles of scientific transparency in uncertainty communication, and suggests that SPELL appropriately flags areas of ecological complexity where expert input remains essential.

SPELL’s handling of ecological complexity shows mixed results. While it successfully captures broad trophic patterns and adapts to regional differences, its treatment of species that span multiple functional roles needs improvement. For instance, the variable classification of anemones and flatfishes between functional groups, while partially reflecting natural ecological flexibility, suggests the need for more nuanced classification approaches. Whilst here we have relied on the LLM’s ‘embedded’ ecological knowledge to classify species, providing additional ecological information from online databases may improve the quality of grouping assignments. Or perhaps, with the rapid rate of LLM capacity improvement, future LLM’s will perform better at ecological tasks such as these. The identification of additional trophic interactions not present in expert matrices (14.5%) requires careful evaluation these could represent either over-connection or potentially valid relationships that merit further investigation.

These validation outcomes suggest SPELL can serve as a useful starting point for ecosystem model development, particularly in its ability to avoid implausible ecological connections and maintain consistent broad-scale trophic structures. However, its outputs require expert review, especially for specialised ecological groups and complex trophic relationships or where there is only a qualitative understanding of ecoystem function. The balance between SPELL’s systematic approach and the need for ecological expertise emerges as a key consideration for its practical application.

### 4.2. Implications for Decision-Making and EBFM

Given the validation outcomes, SPELL shows promise as a rapid prototyping tool for ecosystem-based fisheries management (EBFM). Its ability to avoid spurious ecological connections while maintaining consistent broad-scale trophic structures enables accelerated model development, reducing construction time for model groups and diet matrices from months to hours. However, limitations with specialised groups and single-species functional units mean it should be used as a starting point for expert refinement rather than a standalone solution. In particular, our findings indicate that commercially important species, or species of specific conservation concern, are not always retained as individual functional groups, even when prompts encourage finer resolution. This underscores the need for the implementation of explicit safeguards and verification to ensure management-relevant distinctions are preserved during automated grouping.

To support decision-making, we recommend a hybrid approach that combines AI-driven synthesis with targeted expert input and validation. Specifically, template design will likely benefit greatly from human input to highlight the key groups and species of concern. Further, expert review should be concentrated on outputs flagged by SPELL’s internal uncertainty metrics, such as low diet stability scores or ambiguous groupings, allowing selective triage of high-uncertainty areas. This strategy would enable efficient allocation of expert effort while preserving the benefits of rapid prototyping. SPELL’s systematic uncertainty quantification may also help to identify where additional data collection or expert input is most needed, aligning with principles of uncertainty-aware ecosystem management [20, 31].

This hybrid approach would also address concerns about trust and transparency in AI-assisted modelling. Decision makers often perceive ecological models as ‘black boxes’ with questionable data inputs [3], a concern that may be amplified with AI-based approaches. By integrating expert feedback into the modelling workflow (particularly in areas identified as uncertain), and having an expert assume responsibility for the outputs, SPELL can help mitigate these concerns and foster greater confidence in AI-assisted modelling.

### 4.3. Limitations and Future Development

Our validation results highlight three key limitations of SPELL. First, SPELL’s bias toward generalised classifications, evidenced by its difficulty with single-species functional groups in the GAB comparison, reflects fundamental limitations in how SPELL processes ecological relationships. Second, we tested SPELL with Claude 3.5 Sonnet, a closed-source LLM, which introduces scientific reproducibility challenges. While our validation demonstrates consistent performance, we cannot fully examine the LLM’s decision-making process or potential biases. Third, practical implementation faces computational and data-related constraints. Data harvesting operations proved time-intensive, and SPELL’s performance varied with data availability across regions and ecological roles. Future iterations might benefit from exploring open-source alternatives [27] and developing more transparent decisionmaking processes.

To address the identified limitations, several key areas require further development. First, SPELL’s handling of specialised ecological groups needs improvement, particularly for commercially important single-species units. This could involve developing more sophisticated protocols for identifying and preserving management-relevant distinctions during the grouping process, as different key groupings are more important for fishery management vs. spatial planning, for example. Second, to enhance scientific reproducibility, future versions should explore the capability of other LLMs, including opensource LLMs which can be more transparently assessed than proprietary models like Claude.

Third, systematic validation across diverse ecosystem types is needed to establish operational boundaries. This validation should encompass a range of ecosystems with varying structures, biodiversity levels, and data availability including polar regions, coral reefs, deep ocean habitats, pelagic systems, and upwelling zones and across multiple spatial scales from ocean basins to coastal bays. Testing should pay particular attention to how SPELL handles specialised ecological roles in different contexts.

Fourth, we rely heavily on online databases which are subject to data quality issues and biases. Future work should explore how to incorporate local knowledge and expert judgment into SPELL to address these limitations. This could involve developing more sophisticated data integration methods that combine structured data from online sources with unstructured local knowledge, as well as exploring how to incorporate expert feedback into the AI decision-making process potentially involving weighting the importance of certain sources of data for the LLM. Additionally, it would be valuable to test whether using local or national databases (e.g., Fishes of Australia) alongside global databases can improve SPELL’s accuracy. For example, Australian scientists find that using local data to correct or constrain global database entries substantially improves the quality of other fisheries assessment processes, such as the Ecological Risk Assessment of the Effects of Fishing [21]. Current movements towards FAIR data principles [41] will likely also improve the ability for AI systems to find the most accurate and relevant data sources. Current movements towards FAIR data principles [41] will likely also improve the ability for AI systems to find the most accurate and relevant data sources.

Fifth, future development should incorporate established best practices for food web construction and ecological model building to enhance quality and reliability. Various food web modeling approaches offer methodological standards [8, 19] and diagnostic tools [30] that could strengthen AI-assisted frameworks. More broadly, Good Modelling Practice (GMP) principles [24] emphasize the importance of explicating modelling choices throughout the entire modelling lifecycle to build trust in model insights within their social and political contexts. Integrating these established practices would improve model assessment, facilitate evaluation by management bodies, and help normalize rigorous standards across the modelling community; particularly important as ecological network models increasingly inform resource management decisions.

Sixth, although the grouping prompt instructed the LLM to consider traits such as growth and consumption rates, these were not explicitly validated in this study. Our evaluation focused on ecological roles and diet composition rather than physiological parameters. Future work should assess how well these traits are reflected in functional group assignments, as they are critical for Ecopath dynamics and influence energy flow and biomass turnover. Building on this, SPELL could also be extended to estimate key Ecopath parameters such as production/biomass (PB) and consumption/biomass (QB). These parameters rely on traits already retrieved during data harvesting, and automating their synthesis would further accelerate model parameterization and reduce reliance on manual calculations.

Finally, SPELL’s utility for ecosystem-based fisheries management should be further explored through case studies that evaluate its effectiveness in building models that support management decisions. This could involve comparing the performance of models built using SPELL to those built using traditional methods, as well as assessing SPELL’s ability to support management-relevant analyses such as scenario testing and policy evaluation. While this study focused on marine ecosystems, the underlying framework is not restricted to marine contexts. Its modular design, which integrates global biodiversity databases and user-provided knowledge, can be adapted for freshwater and terrestrial food webs. For example, substituting OBIS with freshwater or terrestrial occurrence databases and adjusting functional group templates would enable application to lakes, rivers, and land-based ecosystems. This flexibility positions SPELL as a broadly applicable tool for accelerating food web model development across diverse ecological domains.

## Acknowledgements

SS was funded by a CSIRO R+ Postdoctoral Fellowship. This project was partially funded by the ‘Futures of Seafood’ project.

## Software Availability

The complete codebase, including all scripts, configuration files, and analysis tools, is available at https://github.com/s-spillias/AI-EwE-Diets. The validation framework, including reference group definitions and classification rules, is documented in the project repository to ensure reproducibility.

## Author Contributions

SS: Conceptualization, Methodology, Software, Validation, Formal analysis, Investigation, Resources, Data Curation, Writing - Original Draft, Writing - Review & Editing, Visualization, Project administration. BF: Validation, Writing - Review & Editing, Supervision, Funding acquisition. FB: Methodology, Software, Validation, Writing - Review & Editing, Supervision. CB: Investigation, Data Curation, Validation, Writing - Review & Editing. JS: Conceptualization, Validation, Investigation, Writing - Review & Editing. RT: Methodology, Software, Validation, Investigation, Writing - Review & Editing, Supervision, Funding acquisition.

## Statement on the Use of Generative AI

Generative AI tools, specifically Claude Sonnet 3.5, were utilized in the preparation of this manuscript to assist with tasks such as language refinement, text structuring, and summarization. All scientific content, data interpretation, and conclusions were independently developed and verified by the authors to ensure accuracy and integrity.

## Supplementary Material

### S1. Data Harvesting Implementation

Our data harvesting system employs DuckDB for efficient querying of PARQUET files, enabling complex joins and aggregations without full memory loading. For species matching across databases, we use structured SQL queries that join on concatenated genus and species names:

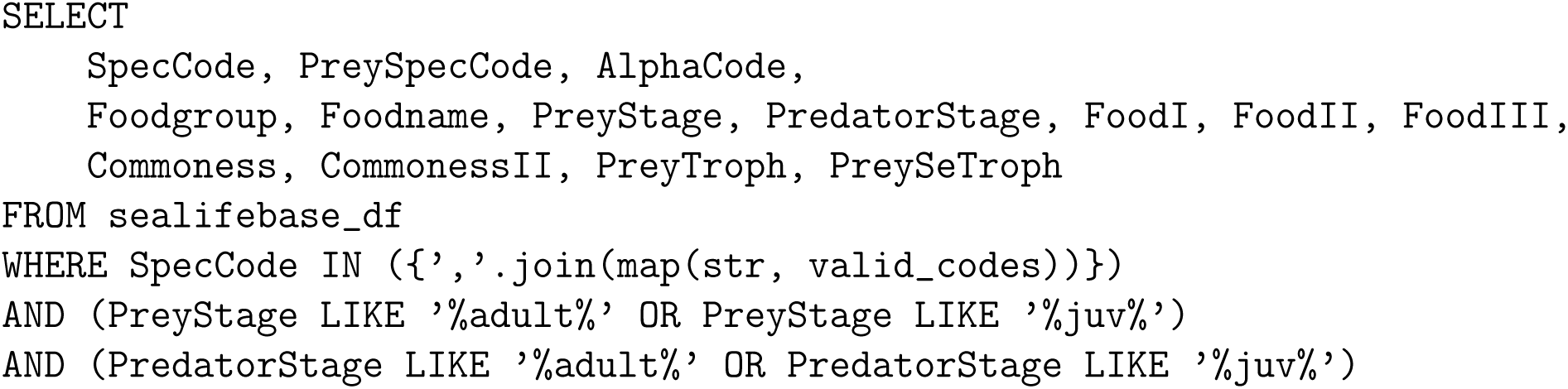

When combining interaction data from GLOBI with diet information, we implement a comprehensive interaction mapping system that creates bidirectional records:

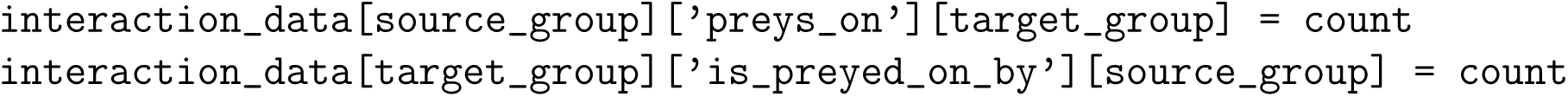

Our data cleaning protocol standardises types by converting numerical values to consistent formats and timestamps to ISO format. We handle null values by removing empty values, ‘NA’ strings, and null entries while preserving data structure. Source tracking maintains database origin information for all data points.

SPELL implements file locking mechanisms for concurrent access, with separate locks for species data and interaction networks. We use exponential backoff retry logic for API interactions, with configurable parameters including maximum retries (5), initial delay (1 second), and maximum delay (60 seconds).

The completion check system verifies the presence of required fields including:

- Complete taxonomic hierarchy
- Species-specific database records (when available)
- Interaction data
- Source attribution
- Data quality indicators

The final JSON output maintains a consistent structure across all species entries, facilitating automated processing in subsequent framework stages.

### S2. Food Web Analysis

**Figure S1:**
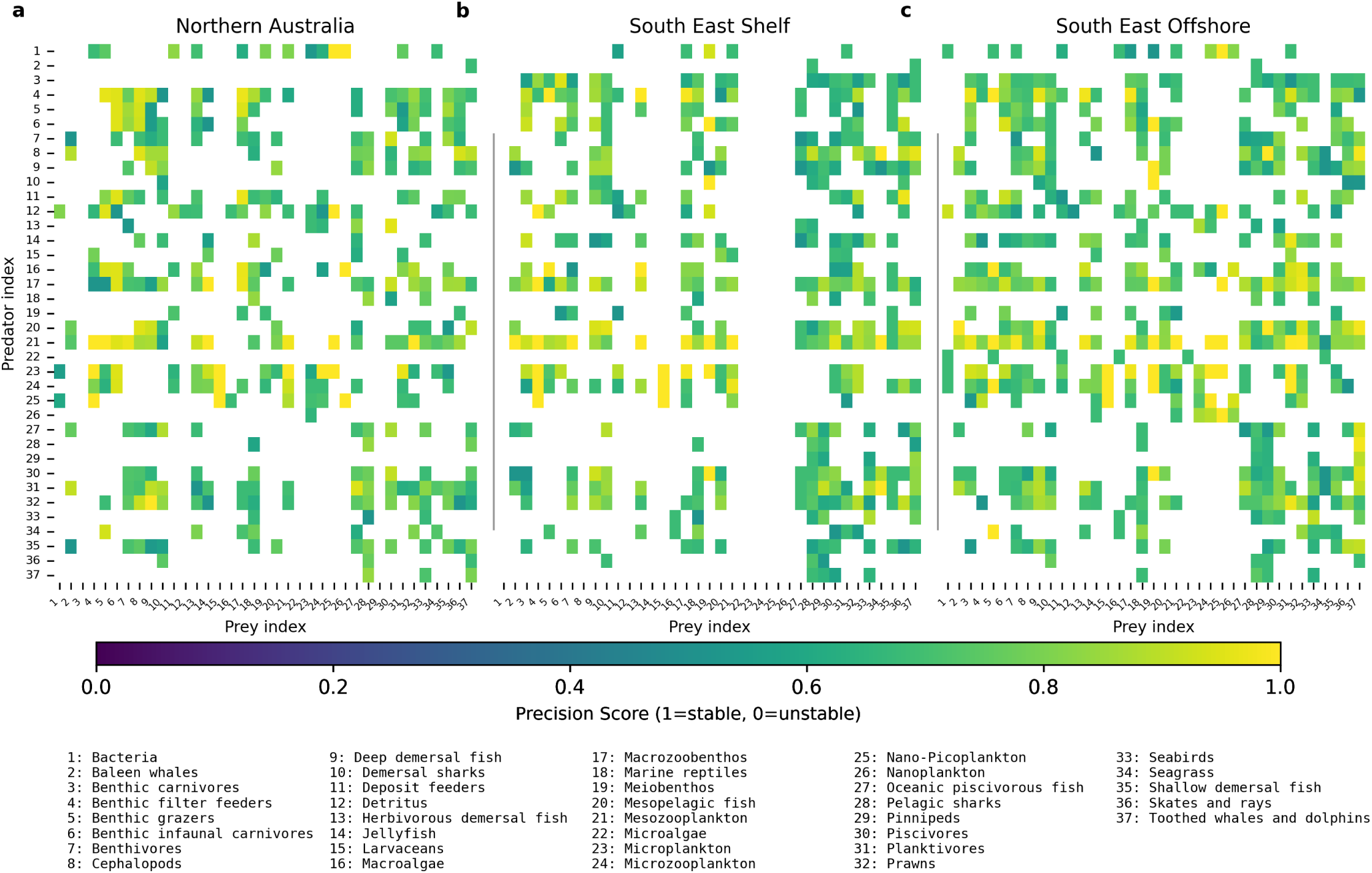
Detailed food web consistency across five iterations for each geographic region. Column names represent predator groups and row names represent their prey groups. Numbers in each cell indicate the mean diet proportions across five iterations, while cell colors indicate the stability score (0-1, where 1.0 = perfect stability; lower values = less stable). White cells represent absent feeding relationships.

**Figure S2:**
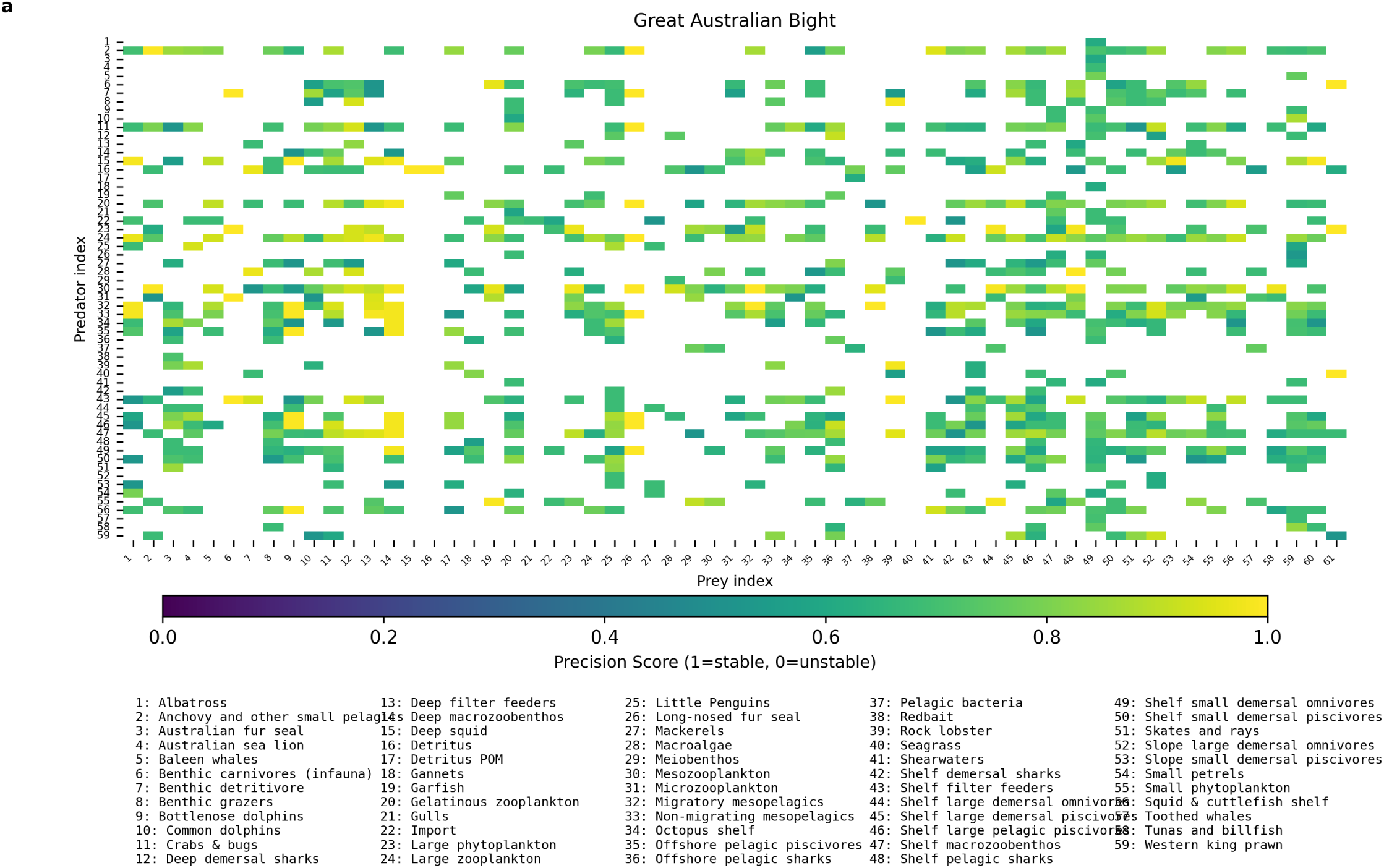
Detailed food web consistency across five iterations for the Great Australian Bight region. Column names represent predator groups and row names represent their prey groups. Numbers in each cell indicate the mean diet proportions across five iterations, while cell colors indicate the stability score (0-1, where 1.0 = perfect stability; lower values = less stable). White cells represent absent feeding relationships.

**Figure S3:**
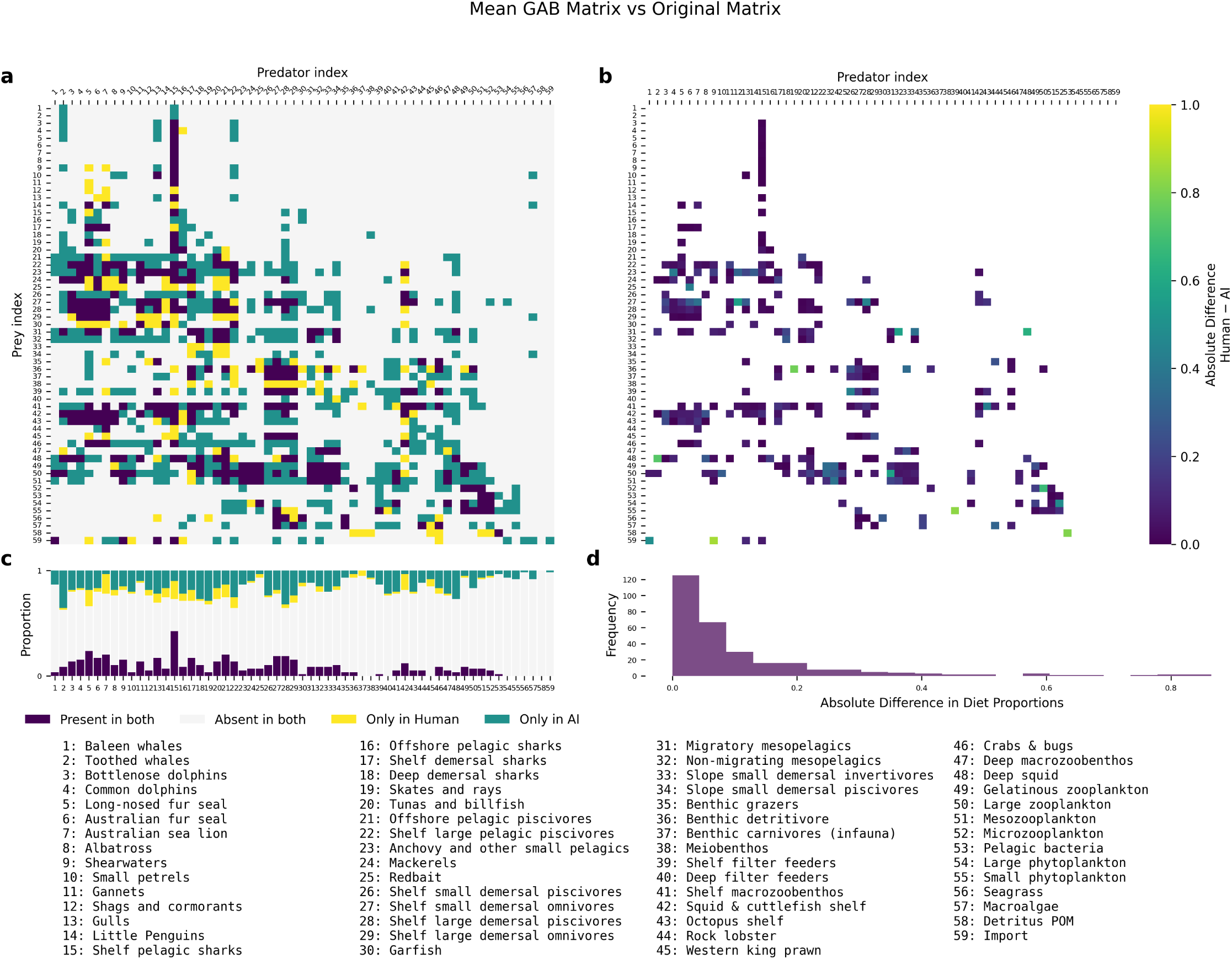
Detailed comparison of food web elements between expert-derived and AI-generated matrices for the Great Australian Bight ecosystem. Panel (a) shows the complete diet matrix with color-coded interaction types. Panel (b) displays the absolute differences in diet proportions between expert and AI matrices where interactions are present in both. Panel (c) shows the proportional breakdown of interaction types for each predator group, while panel (d) presents the frequency distribution of absolute differences in diet proportions.

### S3. Group Stability Analysis

**Figure S4:**
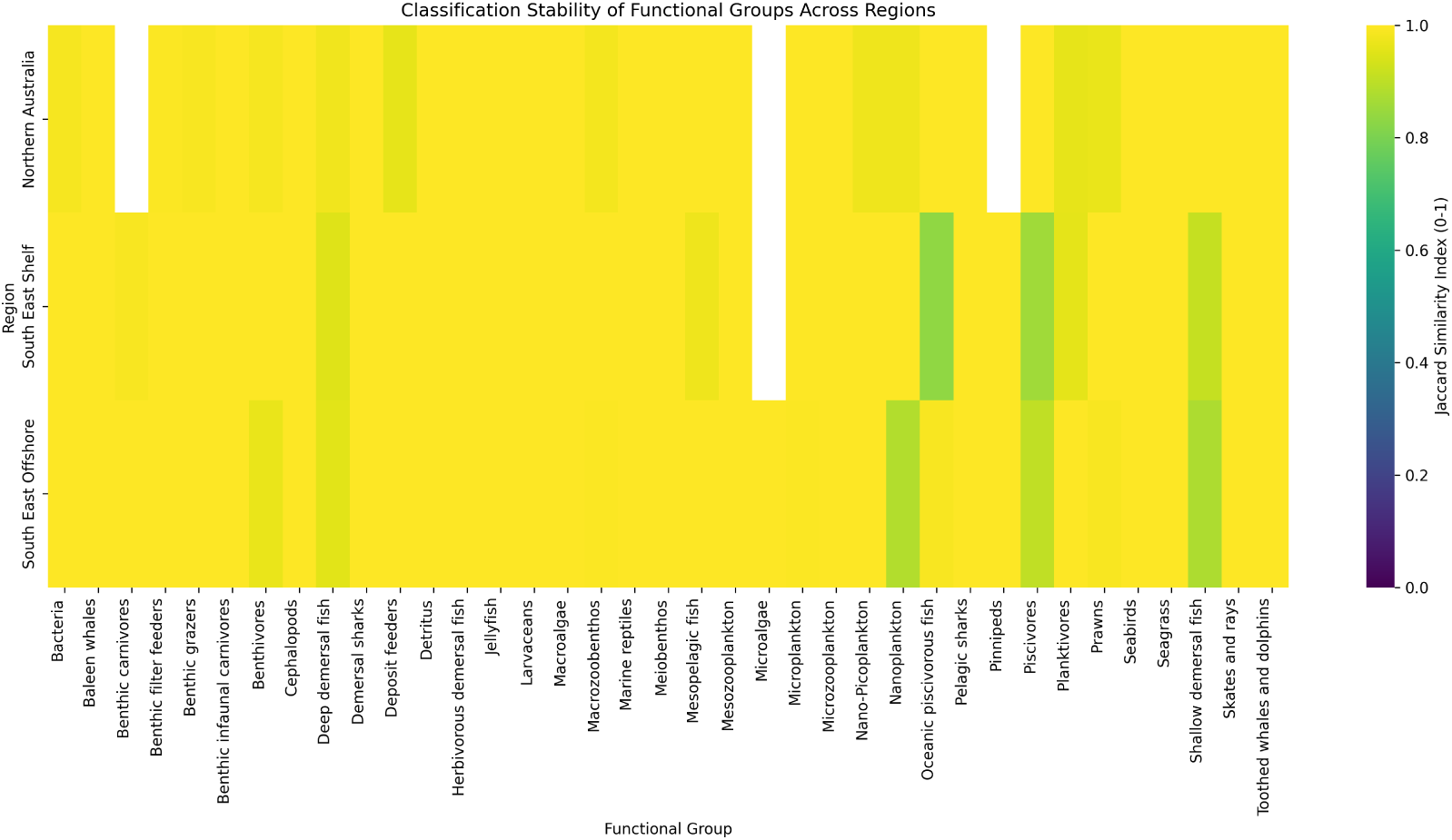
Heatmap showing the stability of functional group classifications across regions. Each cell displays the Jaccard similarity score (ranging from 0.975 to 1.000) between consecutive SPELL iterations, where 1.000 indicates perfect consistency in species assignments. Yellow colors represent higher stability (scores near 1.000), while darker purple colors indicate more variable classifications (scores closer to 0.975). Most functional groups show high stability (>0.99) across all regions, with occasional variations in groups like benthic grazers and deposit feeders, particularly in the Northern Australia region. White indicates groups that were not assigned by the AI system for that region.

### S4. Technical Implementation

#### S4.1. Default Grouping with Descriptions

Table S1 presents the complete template of potential functional groups used by SPELL. This template serves as the initial reference for functional group classification. The LLM uses this template as guidance but may adjust or expand groups during assignment. The final groups in the main text reflect these adjustments.

**Figure S5:**
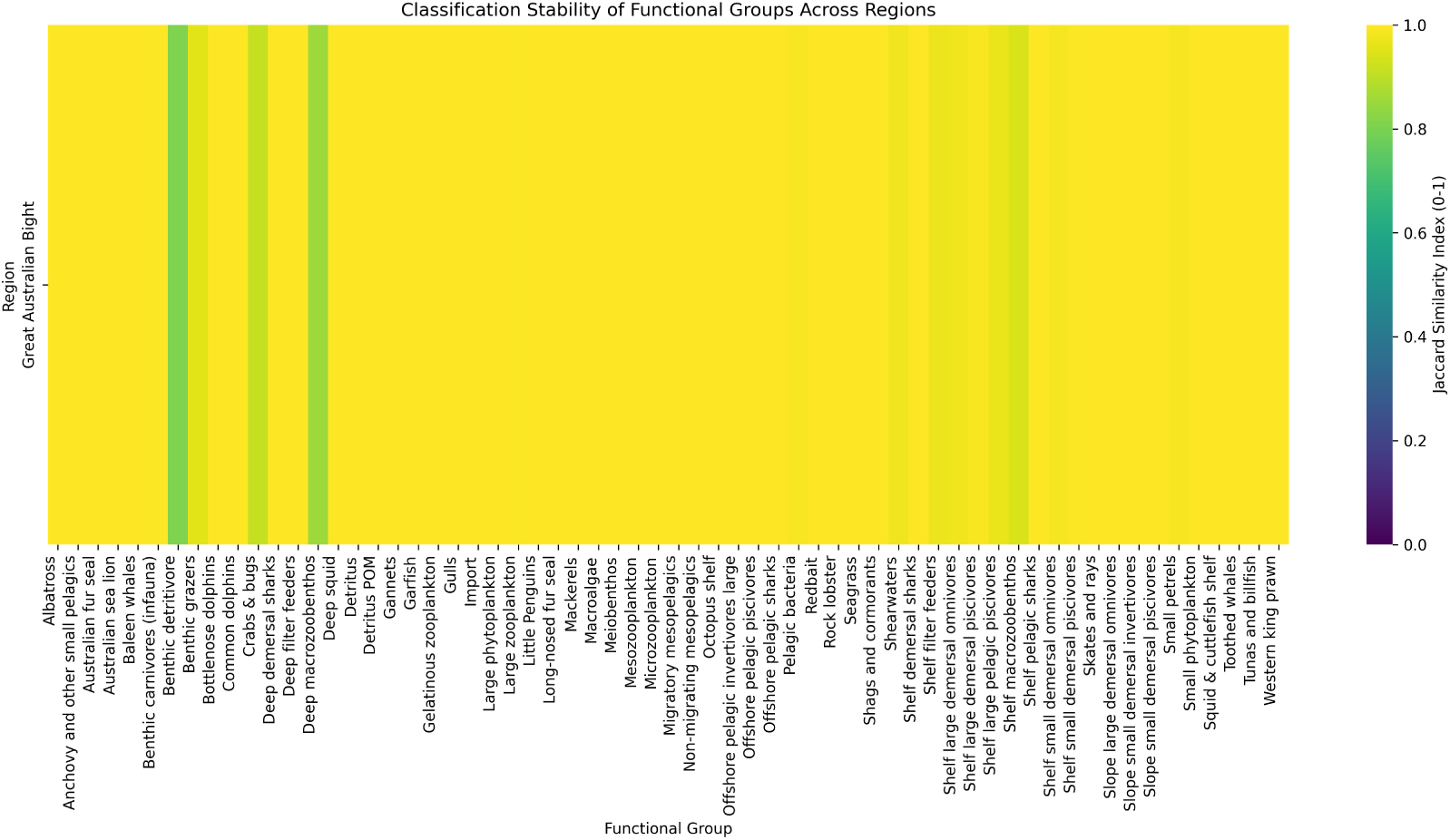
Heatmap showing the stability of functional group classifications across regions. Each cell displays the Jaccard similarity score (ranging from 0.975 to 1.000) between consecutive SPELL iterations, where 1.000 indicates perfect consistency in species assignments. Yellow colors represent higher stability (scores near 1.000), while darker purple colors indicate more variable classifications (scores closer to 0.975). Most functional groups show high stability (>0.99) across all regions, with occasional variations in groups like benthic grazers and deposit feeders, particularly in the Northern Australia region. White indicates groups that were not assigned by the AI system for that region.

**Figure S6:**
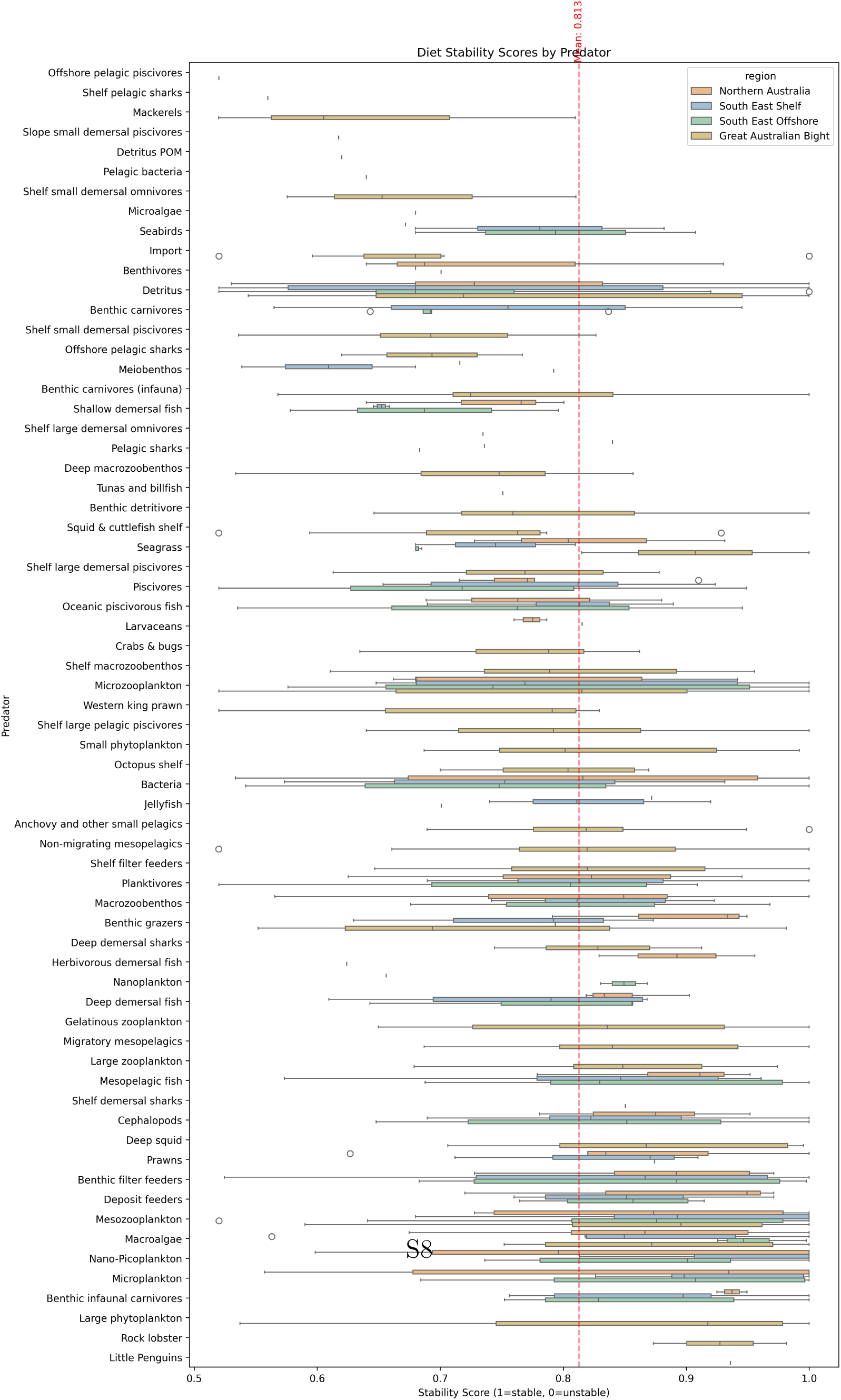
Diet stability scores for substantial interactions (those comprising more than

**Table S1:** Complete Functional Group Template.

| Group Name | Description |
| --- | --- |
| Skates and rays | Bottom-dwelling cartilaginous fish that play a role in controlling benthic prey populations |
| Nearshore and smaller seabirds | Small gulls, terns etc that feed near shore (possibly include penguins here too) - avian predators that link marine and terrestrial ecosystems |
| Albatrosses | Large seabirds that forage exclusively at sea, feeding on marine prey (fishes, squids, gelatinous organisms) |
| Skuas and giant petrels | Large predatory seabirds that feed both at sea and on land, including predation on other birds |
| Fish-eating pinnipeds | Marine mammals (seals, sea lions) that primarily prey on fish in coastal and pelagic ecosystems |
| Invertebrate-eating pinnipeds | Marine mammals (particularly Antarctic seals) that primarily feed on krill and other invertebrates |
| Baleen whales | Large filter-feeding marine mammals that regulate zooplankton populations and contribute to nutrient cycling |
| Orcas | Apex predators that uniquely prey upon other top predators including marine mammals, sharks, and large fish |
| Sperm whales | Deep-diving cetaceans that primarily feed on deep-water squid and fish |
| Small toothed whales and dolphins | Smaller cetaceans that primarily feed on fish and squid in surface and mid-waters |
| Sea snakes | Marine reptiles that prey primarily on fish, particularly eels and fish eggs |
| Crocodiles | Large predatory reptiles in coastal and estuarine waters that prey on fish, birds, and mammals |
| Turtles | Herbivores and omnivores that breed on land |
| Planktivores | Small fishes that feed on plankton, crucial in transferring energy from plankton to larger predators |
| Flying fish | Epipelagic fish capable of gliding above the water surface, important prey for many predators |
| Remoras | Fish that form commensal relationships with larger marine animals, feeding on parasites and food scraps |

Table S1 – Continued
| **Group Name** | **Description** |
| --- | --- |
| Large oceanic piscivorous fish | Fish-eating predators in open ocean environments, mid-sized non-migratory species (e.g. barracuda) |
| Tuna and Billfish | Large oceanic predatory fish, highly mobile, often dive to feed deeper into the water column |
| Shelf small benthivores | Small bodied fish that feed on benthic organisms, playing a key role in benthic-pelagic coupling, live in shelf waters |
| Shelf demersal omnivorous fish | Medium sized demersal fish that feed on invertebrates as well as smaller fish, live in shelf waters |
| Shelf medium demersal piscivores | Medium sized demersal fish living near the bottom in shallow waters, often important in benthic food webs, feed on other fish primarily, live in shelf waters |
| Shelf large piscivores | Fish-eating predatory fishes found in various marine habitats, important in controlling prey fish populations |
| Herbivorous demersal fish | Bottom-associated fish that primarily feed on plants, important in controlling algal growth |
| Slope/deep water benthivores | Small to mid sized fish that feed on benthic organisms and live on the shelf or seamounts |
| Slope/deep demersal omnivorous fish | Medium sized demersal fish that feed on invertebrates as well as smaller fish, live in slope or seamount waters |
| Slope/deep medium demersal piscivores | Medium sized demersal fish that feed on other fish primarily, live in slope or seamount waters |
| Slope/deep large piscivores | Fish-eating predatory fishes found in various marine habitats in deeper water, live in slope or seamount waters |
| Migratory mesopelagic fish | Fish living in the mesopelagic zone, undertake diel vertical migration, important in energy transfer between depths |
| Non-migratory mesopelagic fish | Fish living in the mesopelagic zone, non-migratory species, important in energy transfer between depths |
| Reef sharks | Top predators in coral reef ecosystems, controlling fish populations and maintaining reef health |
| Pelagic sharks | Open-ocean predators that help regulate populations of fishes and squids |

Table S1 – Continued
| Group Name | Description |
| --- | --- |
| Demersal sharks | Bottom-dwelling sharks, including dogfishes, that control populations of fishes and invertebrates on and near the seafloor |
| Cephalopods | Intelligent mollusks like squid and octopus, important predators in many marine ecosystems |
| Hard corals | Reef-building colonial animals that create complex habitat structure through calcium carbonate deposition |
| Soft corals | Colonial animals that contribute to reef habitat complexity without building calcium carbonate structures |
| Sea anemones | Predatory anthozoans that can form symbiotic relationships with fish and crustaceans |
| Hydrothermal vent communities | Specialized organisms living around deep-sea vents, including chemosynthetic bacteria and associated fauna |
| Cold seep communities | Organisms adapted to methane and sulfide-rich environments on the seafloor |
| Deep-sea glass sponges | Filter-feeding animals that create complex deep-water habitats and are important in silicon cycling |
| Sea cucumbers | Deposit-feeding echinoderms important in sediment processing and bioturbation |
| Sea urchins | Herbivorous echinoderms that can control macroalgal abundance and affect reef structure |
| Crown-of-thorns starfish | Coral-eating sea stars that can significantly impact reef health during population outbreaks |
| Benthic filter feeders | Bottom-dwelling organisms that filter water for food, important in nutrient cycling and regulating water quality in various depths - bivalves, crinoids, sponges |
| Macrozoobenthos | Mobile large bottom-dwelling invertebrates in both shallow and deep waters, important in benthic food webs and bioturbation (predatory or omnivorous) |
| Benthic grazers | Bottom-dwelling organisms that graze on algae and detritus, influencing benthic community structure |
| Prawns | Small crustaceans that are important in benthic and pelagic food webs |

Table S1 – Continued
| Group Name | Description |
| --- | --- |
| Meiobenthos | Tiny bottom-dwelling organisms, important in sediment processes and as food for larger animals |
| Deposit feeders | Animals that feed on organic matter in sediments, important in nutrient cycling |
| Benthic infaunal carnivores | Predatory animals living within the seafloor sediments |
| Sedimentary Bacteria | Microscopic organisms crucial in nutrient cycling and the microbial loop in marine ecosystems |
| Large carnivorous zooplankton | Fish larvae, arrow worms and other large predatory zooplankton |
| Antarctic krill | Key species in Antarctic food webs, particularly important as prey for whales, seals, and seabirds |
| Ice-associated algae | Microalgae living within and on the underside of sea ice, important primary producers in polar regions |
| Ice-associated fauna | Specialized invertebrates living in association with sea ice, important in polar food webs |
| Mesozooplankton | Medium-sized zooplankton (200 $\mu\text{m}$ to 2 cm) that feed on smaller plankton and serve as food for larger animals |
| Microzooplankton | Tiny zooplankton (20 $\mu\text{m}$ to 200 $\mu\text{m}$) that graze on phytoplankton and bacteria, forming a crucial link in the microbial food web |
| Pelagic tunicates | Including larvaceans, salps, and pyrosomes, important in marine snow formation and carbon cycling |
| Jellyfish | Predatory gelatinous species |
| Diatoms | Larger phytoplankton (20 $\mu\text{m}$ to 200 $\mu\text{m}$), silica dependent important primary producers in marine ecosystems |
| Dinoflagellates | Mixotrophic species (20 $\mu\text{m}$ to 200 $\mu\text{m}$) that can switch between primary production and consumption as needed |
| Nanoplankton | Plankton ranging from 2 $\mu\text{m}$ to 20 $\mu\text{m}$ in size, including small algae and protozoans |
| Picoplankton | Plankton ranging from 0.2 $\mu\text{m}$ to 2 $\mu\text{m}$ in size, including both photosynthetic and heterotrophic organisms |

Table S1 – Continued
| Group Name | Description |
| --- | --- |
| Microalgae (microphytobenthos) | Microscopic algae that live on the seafloor or attached to other organisms |
| Pelagic bacteria | Watercolumn dwelling bacteria, consume marine snow amongst other things |
| Seagrass | Marine flowering plants that form important coastal habitats and nursery areas |
| Mangroves | Salt-tolerant trees forming critical coastal nursery habitats and protecting shorelines |
| Salt marsh plants | Coastal vegetation adapted to periodic flooding, important in nutrient cycling and shoreline protection |
| Macroalgae | Seaweeds of various sizes that provide habitat and food for many species, including both canopy and understory forms |
| Symbiotic zooxanthellae | Photosynthetic dinoflagellates living within coral and other marine invertebrates |
| Cleaner fish and shrimp | Species that remove parasites from other marine animals, important in reef health |
| Discards | Carrion and freshly discarded material from fisheries activities |
| Detritus | Labile components of natural death and waste |

#### S4.2. Retrieval-Augmented Generation Implementation

We implement a retrieval-augmented generation system using ChromaDB for vector storage and document management. Document processing begins with LlamaParse conversion of source materials to markdown format, preserving structural elements while enabling consistent text extraction across document types. We segment documents using a token-aware chunking strategy with a 2000-token maximum size, determined through empirical testing to balance context preservation with model limitations.

Document processing follows a two-phase approach. The initial phase generates embeddings for each document chunk using Azure OpenAI’s textembedding-3-small model, storing them in ChromaDB’s PersistentClient. SPELL maintains an indexed_files.json registry to track processed documents. The second phase handles incremental updates, identifying and processing only new content when documents are added to the source directory.

For diet composition analysis, we implement a two-stage query process. The first stage employs a simple query to retrieve relevant document chunks:

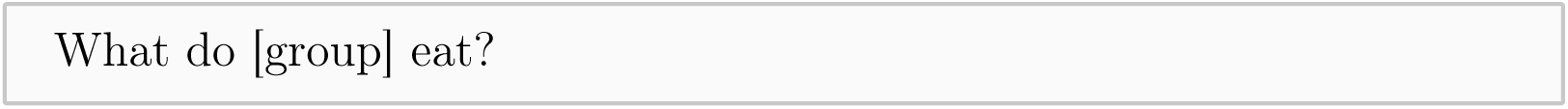

SPELL embeds this query using the same Azure OpenAI model and performs vector similarity search to identify relevant document chunks. These results combine with structured data sources including species occurrence frequencies, food category classifications, and GLOBI interaction data to form a comprehensive input for the second stage.

We implement comprehensive error handling throughout the pipeline. SPELL employs exponential backoff retry logic for API interactions, with configurable parameters including maximum retries (10), initial delay (1 second), and maximum delay (300 seconds). For model interactions, we utilise LlamaIndex’s query engine with zero-temperature sampling to ensure deterministic responses. SPELL supports multiple language model backends, enabling flexible deployment based on availability and performance requirements.

SPELL maintains separate storage contexts for different document collections through ChromaDB’s collection management. This separation prevents cross-contamination between knowledge bases while enabling efficient parallel processing. We track document citations throughout the retrieval process, maintaining provenance information for all retrieved content. The complete implementation, including embedding generation, chunking algorithms, and query processing functions, is available in the project repository.

## Notes

https://github.com/s-spillias/AI-EwE-Diets

### Competing Interest Statement

The authors have declared no competing interest.

### Summary of Updates

This revisions is the result of responding to peer review comments. Significant changes include: 1) Renaming of the framework Synthesising Parameters for Ecosystem modelling with LLMs (SPELL) for ease of communication. 2) Clarification of methods and results 3) Revising figures for clarity

